# Convergent neural representations of acute nociceptive pain in healthy volunteers: A large-scale fMRI meta-analysis

**DOI:** 10.1101/779280

**Authors:** Anna Xu, Bart Larsen, Erica B. Baller, J. Cobb Scott, Vaishnavi Sharma, Azeez Adebimpe, Allan I. Basbaum, Robert H. Dworkin, Robert R. Edwards, Clifford J. Woolf, Simon B. Eickhoff, Claudia R. Eickhoff, Theodore D. Satterthwaite

## Abstract

Characterizing a reliable, pain-related neural signature is critical for translational applications. Many prior fMRI studies have examined acute pain-related brain activation in healthy participants. However, synthesizing these data to identify convergent patterns of activation can be challenging due to the heterogeneity of experimental designs and samples. To address this challenge, we conducted a comprehensive meta-analysis of fMRI studies of stimulus-induced pain in healthy participants. Following pre-registration, two independent reviewers evaluated 4,927 abstracts returned from a search of 8 databases, with 222 fMRI experiments meeting inclusion criteria. We analyzed these experiments using Activation Likelihood Estimation with rigorous type I error control (voxel height *p* < 0.001, cluster *p* < 0.05 FWE-corrected) and found a convergent, largely bilateral pattern of pain-related activation in the secondary somatosensory cortex, insula, midcingulate cortex, and thalamus. Notably, these regions were consistently recruited regardless of stimulation technique, location of induction, and participant sex. These findings suggest a highly-conserved core set of pain-related brain areas, encouraging applications as a biomarker for novel therapeutics targeting acute pain.

**HIGHLIGHTS:**

- Pain stimulation recruits a core set of pain-related brain regions.
- This core set includes thalamus, SII, insula and mid-cingulate cortex.
- These regions were recruited regardless of stimulus modality and stimulus location.

## 1. INTRODUCTION

Chronic and acute pain are global medical issues affecting at least 20% of adults globally and they are often accompanied by both comorbid psychological disorders and significant disability in daily activities (Goldberg & McGee, 2011). Given the prevalence of pain conditions, there is a need to develop tools capable of translating the subjective report of pain into an objective measure (i.e., a pain biomarker) that can be used in the development of novel treatments. Recently, neuroimaging approaches have been used to examine pain-related brain activity as a physiological biomarker of pain for treatment development (Cowen et al., 2015; Labus et al., 2015; Wager et al., 2013; Woo et al., 2017). Though the challenge of pain biomarker development is increased when chronic pain conditions are analyzed, a growing body of work examining the neural correlates of experimentally-induced, nociceptive pain in healthy volunteers has led to important insights into the mechanisms and characteristics of how the sensation of pain arises, including its cognitive, affective, and sensory dimensions that may not be reflected in self-report scales (Apkarian et al., 2005; Hofbauer et al., 2001; Robinson et al., 2013; Talbot et al., 1991; Tracey & Mantyh, 2007; Treede et al., 1999; Woo & Wager, 2016). While this work has been critical in elucidating the circuits recruited by acute nociceptive pain under a variety of experimental contexts, synthesizing these data to identify convergent patterns of pain-related brain activation can be challenging due to heterogeneity of experimental designs and samples.

To integrate findings across neuroimaging experiments, meta-analysis provides a powerful approach to quantitatively identify consistent brain regions activated during pain (Wager et al., 2009). Prior meta-analyses have identified several brain regions that are engaged under pain conditions, including the secondary somatosensory cortex (SII), insula, cingulate cortex, and thalamus. Other regions have been identified somewhat less reliably, including primary somatosensory cortex (SI), striatum, cerebellum, supplementary motor area (SMA), primary motor area (M1), periaqueductal gray (PAG), prefrontal cortex (PFC), certain areas in parietal cortices, and the parahippocampal gyrus (Apkarian et al., 2005; Duerden & Albanese, 2013; Farrell et al., 2005; Jensen et al., 2016; Lanz et al., 2011; Peyron et al., 2000; Tanasescu et al., 2016). Subsequent meta-analyses have sought to parse this pain network further by investigating neural responses specific to different pain induction modalities, such as thermal pain (Farrell et al., 2005; Friebel et al., 2011; Jensen et al., 2016), and to different stimulation location (Duerden & Albanese, 2013; Jensen et al., 2016; Lanz et al., 2011). These meta-analyses have provided a substantial advance in our understanding of the brain’s pain network. However, the significant increase in functional magnetic resonance imaging (fMRI) experiments in pain since the most recent meta-analyses (Jensen et al., 2016; Tanasescu et al., 2016) as well as an influx of articles standardizing rigorous procedures for meta-analyses (Eickhoff et al., 2016; Müller et al., 2018) suggest that an update is warranted. Furthermore, based on recent evidence of inadequate Type I error control using previously-typical statistical methods, we considered it worthwhile to revisit the topic of identifying brain regions consistently recruited by diverse pain-inducing stimuli while adhering to contemporary standards in the field (Eklund et al., 2016; Müller et al., 2018).

Accordingly, we conducted a comprehensive meta-analysis of fMRI studies of experimentally-induced pain in healthy volunteers; the analysis included findings from 222 experiments. We first sought to replicate and extend the findings from previous meta-analyses, incorporating more recent studies in this rapidly moving field. Second, we applied standards for Type I error correction to our analyses according to current standards in the field. Third, we assessed differences in pain responsiveness associated with differences in pain stimulation modality (thermal, electrical, mechanical, or chemical), location of stimulation (visceral or somatic, left or right side of body, proximal or distal extremity), and sample composition (participant sex). As described below, our results revealed highly convergent evidence for the existence of a core set of brain regions associated with acute nociceptive pain in healthy participants. This core set was present across different samples and experimental designs, encouraging its use as biomarker of acute pain that could be useful for experimental therapeutics.

## 2. METHODS

Our methodology adheres to PRISMA and field-standard guidelines for meta-analyses (Moher et al., 2009; Müller et al., 2018). Following pre-registration, we first performed a literature search for fMRI experiments of experimentally induced pain in healthy participants using eight databases and then searched for references in reviews identified in the database. Titles and abstracts returned by this search were first evaluated for full-text screening. Full text articles were evaluated to see if they met defined inclusion criteria (see **Figure 1** for PRISMA chart detailing screening process). This screening process resulted in a total of 222 fMRI experiments from 200 articles that were included in this study. See specific details below. Coordinate data from these experiments were then extracted and analyzed using activation likelihood estimation (ALE).

**Figure 1.**
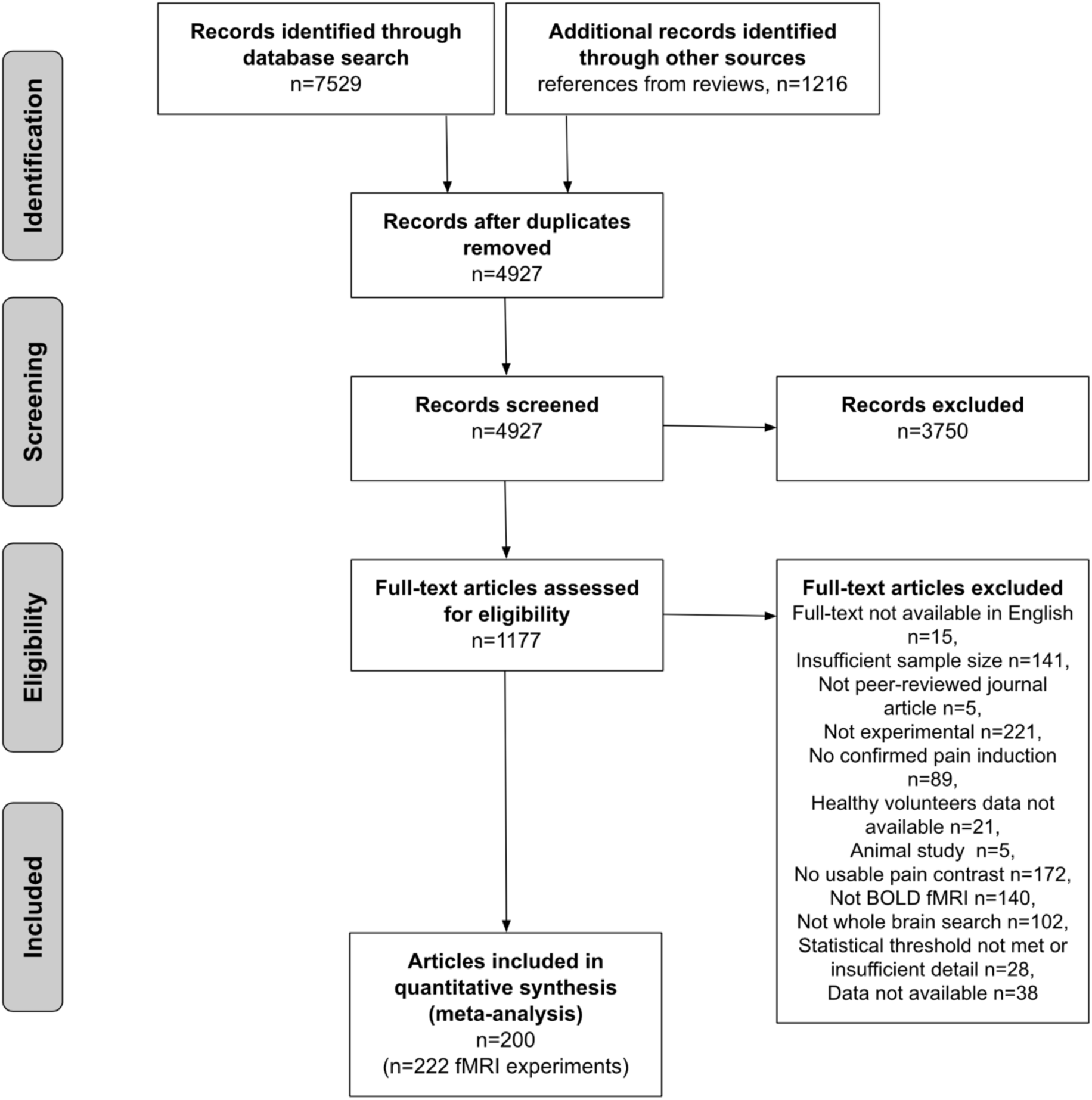
PRISMA flowchart for study inclusion.

### 2.1. Literature search

We performed a literature search for fMRI experiments of experimentally induced pain in healthy volunteers using both database searches and references cited in review articles, meta-analyses, and component studies. The final literature search took place on August 1, 2018 and was restricted to articles published from 1990 to August 1, 2018.

#### 2.1.1. Database search

The following standard literature databases were searched: PUBMED/MEDLINE, EMBASE, Web of Science, Cochrane Library, and PsycINFO. We used the following search terms: (“MRI” or “magnetic resonance imaging” or “fMRI” or “BOLD” or “brain mapping”) AND (“pain” or “noxious” or “nociception”). Additional inclusion criteria consisted of experiments conducted in humans, publications that were in English, and articles that appeared in peer-reviewed journals (e.g., not conference papers).

We supplemented our database search with the following existing fMRI data repositories: NeuroSynth (Yarkoni et al., 2011), brainspell (Toro, 2015), and BrainMap (Laird et al., 2005). In these repositories, we searched for records using the keyword “pain.”

#### 2.1.2. Reference search from reviews

To identify potential candidate studies from reference lists, we also screened the resulting abstracts for editorials, review articles, and meta-analyses related to pain. In editorials (i.e., non-systematic reviews), we identified titles in reference sections that seemed likely to include a pain experiment. In systematic reviews, we considered all articles that the authors identified for inclusion. If a systematic review included a meta-analysis of experimentally induced pain in healthy volunteers (e.g., Friebel et al., 2011; Duerden et al., 2013; Jensen et al., 2016; Tanasescu et al., 2016), we automatically included all studies within the meta-analysis to be screened for full text.

The database search yielded a total of 7,529 articles. We additionally compiled references from existing review articles which yielded a total of 1,216 articles (total of 8,754 records). After removing duplicates, we screened a total of 4,927 abstracts.

### 2.2 Inclusion Criteria

Experiments were only included in the meta-analysis if they contained a within-subject “pain > baseline” contrast (e.g., “pain > rest”, “pain > innocuous stimuli” or parametric modulation of pain) that was not confounded by other experimental manipulations that could impact the acute nociceptive pain induction (e.g., treatment manipulations prior to the pain induction, such as drug infusions or placebo). If at least one experiment in an article satisfied this initial requirement, it was evaluated according to the following inclusion criteria:

1. The experiment was from a peer-reviewed journal article written in English.
2. The experiment considered healthy, human participants over the age of 18. To satisfy this criterion, the study must explicitly report that all participants were healthy or free of medical or psychiatric disorders.
3. The experiment included at least 10 participants.
4. The experiment induced physical pain that was confirmed to be painful by participants. Confirmation of experienced pain could be in the form of an explicit report of the induction being painful, participant ratings of experienced pain during the scan session, or the use of a pain stimulus that was titrated to a threshold pre-determined to be painful by participants in the experiment.
5. Brain responses to induced pain were monitored using fMRI.
6. The field of view and reported results included the whole brain (i.e., region of interest analyses were excluded). This criterion was imposed so as to prevent bias towards *a priori* regions putatively thought to be involved in pain.
7. The experiment reported results in a standard stereotaxic reference space coordinate system (MNI or Talairach space).
8. Results met current statistical standards for conventional cluster identification. Specifically, we only included experiments that reported activation at a voxel-level threshold of *p* < 0.001 (uncorrected) or a corrected cluster probability of *p* < 0.05. We also excluded experiments that did not report their methods and results in sufficient detail to conclude whether they met our statistical threshold criteria.

Additionally, if experiments in articles did not report relevant results but met the inclusion criteria, we e-mailed the corresponding authors and included the experiment if data was provided.

#### 2.2.2. Abstract and Full Text Assessment

Two independent reviewers (AX, EBB) confirmed the inclusion or exclusion of each abstract for full text screening. Abstracts were first assessed as to whether they included a physical pain contrast in healthy volunteers and whether they measured task-based blood-oxygen-level-dependent (BOLD) responses. In this stage of screening, we only excluded abstracts that explicitly mentioned (1) having a sample size of less than 10 subjects; (2) using only a neuroimaging modality that was not fMRI, such as EEG; or (3) only including animal experiments. Note that in this stage of screening, we did not exclude any papers that involved a clinical population, used resting-state fMRI, or involved a treatment or intervention. These criteria allowed us to assess parts of seemingly irrelevant papers that may have included relevant experiments for analyses, such as including a healthy subsample (e.g., the control sample) or a task-based measure involving acute nociceptive pain (e.g., pain inductions either pre-treatment or post-resting-state). Full text articles from included abstracts were then assessed for whether they met our inclusion criteria (see section 2.2.1. Inclusion Criteria). Finally, at least two independent reviewers confirmed the decision for inclusion of articles marked for inclusion in the final analysis (AX, BL, EB). In cases of reviewer decision disagreement, a senior third reviewer (TS) evaluated the article.

### 2.3. Data extraction

Coordinates and information about each experiment were extracted manually by at least one author (AX or VS) and checked independently by another member of the study team (AX or VS). The following information about each paper was extracted: sample size; whether the coordinate space was MNI or Talairach; modality of pain stimulus (e.g., thermal, electrical, mechanical, or chemical); side the stimulus was induced; whether the stimulus was on the arm (not including hand), leg (not including foot), hand, or foot; whether the pain stimulus was visceral (e.g., esophageal distension); and whether the reported activation included a non-painful control stimulus (e.g. painful > innocuous contrast).

In cases where studies contained multiple relevant pain contrast results from a single experiment, we chose to use results that were most likely to demonstrate a clear nociceptive pain-specific signal. For example, when experiments contained separate analyses based on the intensity of the pain induction (e.g., one contrast for moderate pain and another for high pain), we chose the contrast for the highest intensity of pain reported (e.g., high pain). For experiments with separate analyses based on a subjective rating of pain and based on an objective intensity, we included results based on the objective intensity rather than the subjective rating.

If multiple experimental contrasts included in a single article could be used for different sets of meta-analyses (see section 2.4.2.), we first pooled coordinates from each experiment into one set of coordinates for that particular article and treated this set of coordinates as one experiment in our primary meta-analysis (Turkeltaub et al., 2012). This approach ensured that we only used one set of coordinates per article, so that one specific article could not be weighted more heavily than others due to the presence of multiple relevant experiments (Müller et al., 2018; Turkeltaub et al., 2012). For example, for our primary meta-analysis of experimentally-induced pain, if an article contained multiple experiments using different types of pain stimulation (e.g., thermal and mechanical), we pooled the results from both types of pain stimulation together and treated the results as if they were derived from one experiment. For any additional analyses (see section 2.4.2.), we evaluated these contrasts separately(e.g., the thermal set of coordinates would be tagged as thermal pain and treated as one experiment while the mechanical set of coordinates would be tagged as mechanical pain and treated as another experiment).

### 2.4. Coordinate based meta-analysis

#### 2.4.1. Activation likelihood estimation (ALE)

We conducted meta-analyses using the coordinate-based meta-analytic method activation likelihood estimation (ALE) (Turkeltaub et al., 2002) using the revised algorithm that allows for random effects inference (Eickhoff et al., 2009; Eickhoff et al., 2012; Rottschy et al., 2012). The main effect for a particular condition of interest is defined by the convergence of activation from all relevant experiments included in analyses.

Briefly, for each experiment included, ALE treats coordinates for the foci of reported clusters as the center of an uncertainty function modeled by a 3D Gaussian probability distribution. The full width at half-maximum (FWHM) of this 3D Gaussian kernel was determined by empirical data on between-subject and between-template (i.e., MNI or Talairach space coordinates) variance. Specifically, the algorithm takes into account between-subject variance by using a tighter Gaussian distribution for experiments with greater sample sizes to represent that these experiments should provide more reliable results of a true activation effect (Eickhoff et al., 2009), and it takes into account between-template differences by transforming coordinates reported in Talairach coordinates into MNI coordinates (Lancaster et al., 2007). This model then provides probabilities for all activation foci in each experiment, which were combined for each voxel, resulting in an individual modeled activation (MA) map for each experiment. By taking the union across all the MA-maps, we generated voxelwise ALE scores that describe the convergence of results at each particular location (Eickhoff et al., 2009). Note that MA-values reflect data for a single experiment while ALE-values integrate data across multiple experiments.

For ALE maps, the *p*-value was defined as the proportion of values obtained under a null distribution reflecting a random spatial association between experiments. The resulting non-parametric *p* values were subsequently thresholded using voxel height threshold of *p* < 0.001, reflecting current recommendations for best practices (Eklund et al., 2016). At this voxel height, the significance of cluster extent was estimated using 10,000 Monte-Carlo simulations; this distribution was calculated specifically for each meta-analysis conducted (Eickhoff et al., 2012; Rottschy et al., 2012). Clusters were considered significant if they achieved a family-wise corrected significance of *p* < 0.05. Prior to display, *p*-values were then transformed into *z* scores. All results were labeled using either SPM Anatomy Toolbox v2.2 (Eickhoff et al., 2005) or Harvard-Oxford Structural Atlas (Kennedy et al., 1998) distributed by FSL. Thalamic parcellations included with SPM Anatomy Toolbox were based on the Thalamic Connectivity Atlas (Behrens et al., 2003).

##### 2.4.1.1. Statistical contrasts

To analyze differences in convergent activation (i.e., differing convergence of results) between two different groups of experiments, we computed the voxel-wise differences between the cluster-level FWE-corrected maps derived from the individual main effect analyses (as described above). To determine the significant difference in ALE scores, we first generated a null distribution of ALE-score differences by randomly permuting the labels of all experiments, dividing them into two groups of the same sizes as the original analysis, and calculating the ALE-scores for these two randomly permuted groups for all voxels in the brain. We repeated this process 10,000 times and tested the observed differences in ALE-scores against the derived null distribution. We thresholded probability values at *p* < 0.001 and inclusively masked them by the main effects for the particular condition of interest. Finally, we applied an extent-threshold of *k* > 25 voxels. It is important to note, however, that an unequal proportion of experiments in the two groups of interest could bias the observed results. To address this concern, we conducted chi-squared tests for differences in proportion of experiments between the two groups of interest in each of our planned contrasts, and we limited our analyses to comparisons that did not significantly differ in number of experiments (at *p* < 0.01).

##### 2.4.1.2. Conjunction Analyses

To analyze voxels where a significant effect was present in two different groups of experiments, we computed their conjunction using the conservative minimum statistic (Nichols et al., 2005). This approach is equivalent to identifying the intersection between each of the cluster-level FWE-corrected maps of the main effects for the two groups of experiments (Caspers et al., 2010). We then applied an extent-threshold of *k* > 25 voxels to exclude smaller regions of presumably incidental overlap between the two maps of the main effects.

#### 2.4.2. Effects of interest

We primarily focused on finding areas consistently reported to be activated in response to noxious stimuli inducing acute pain. We first assessed areas converging in activation in response to pain using all of our extracted coordinate data (with only one set of coordinates per article). However, because some of our included experiments used a contrast of “pain > rest” while others used a contrast of “pain > innocuous stimuli”, we conducted secondary analyses examining the main effect of the experiments using the contrast “pain > rest” and then the experiments using “pain > innocuous stimuli.” A contrast between experiments using “pain > rest” contrast and “pain > innocuous stimuli” contrast, however, was not computed due to significant differences in the proportion of experiments using the “pain > rest” contrast and the “pain > innocuous stimuli” contrast.

Next, given the heterogeneity of the different pain induction techniques used in the included experiments, we conducted additional contrast and conjunction analyses that examined the effect of different modalities and locations of pain inductions as well as the sex of the sample. The following analyses were conducted:

- To analyze the effect of pain modalities considered, we examined experiments inducing thermal pain (e.g., heat, cold), mechanical pain (e.g., pin prick, pressure, distension), electrical pain (e.g., electrical stimulation), and chemical pain (e.g., capsaicin). To explicitly compare different modalities, we conducted between-experiment contrasts comparing thermal and non-thermal experiments as well as electrical and mechanical experiments. Further analyses comparing other modalities could not be reliably conducted based on significant differences in the proportion of experiments available.
- To examine effects of the location of induced pain, we conducted three separate contrasts. First, we evaluated laterality by contrasting experiments where pain was induced on the left side versus the right side of the body. Second, we examined differences in the effect of inducing pain on the extremities at proximal (i.e., the arm or leg) versus distal (i.e., the hand or foot) locations, which have different densities of nociceptors. Third, we examined differences in brain activation for visceral (e.g., rectal distension) versus non-visceral (i.e., somatic) pain. Notably, we only included non-visceral mechanical pain, because all visceral pain inductions were mechanical.
- Finally, by comparing experiments that included *only* male or *only* female participants, we sought to evaluate sex differences in the pain response (e.g., Coen et al., 2008).

#### 2.4.5. Post-hoc diagnostics

To address bias and heterogeneity of experiments included in our primary meta-analysis of pain, we calculated the contribution of each experiment by computing the ratio of ALE-scores of all voxels in a specific cluster with and without each experiment. Similarly, we calculated the contribution of different conditions of interest from groups of experiments (e.g., thermal, mechanical, right-sided pain, left-sided pain, etc.). These analyses provided an estimate of how the ALE-score changed when the experiment or group of experiments in question was removed (see Cieslik et al., 2016 for example). However, given the large number of experiments included in this study, undue influence of a single experiment was relatively unlikely (Eickhoff et al., 2012).

## 3. RESULTS

Our search, screening, and evaluation yielded a total of 222 experiments from 200 articles that met inclusion criteria as confirmed by two independent reviewers (**Figure 1**). Of these 222 experiments, we meta-analyzed 200 experiments for the main effect of induction of a reported sensation of pain. Among these, 62 experiments used a “pain > innocuous” contrast and 134 experiments used a “pain > rest” contrast. The remaining 3 experiments examined a parametric modulation of pain.

For modality-specific analyses, we meta-analyzed 107 thermal pain experiments (inducing heat pain or cold pain) and 98 non-thermal pain experiments. The non-thermal pain experiments included 39 experiments inducing electrical pain (e.g., electric shocks), 46 inducing mechanical pain (e.g., pressure pain, distension), and 13 inducing chemical pain (e.g., capsaicin). For location-specific analyses, we meta-analyzed 92 left-sided pain experiments, 66 right-sided pain experiments, 68 experiments inducing pain in distal extremities, 85 experiments inducing pain in proximal extremities, 17 experiments inducing visceral pain, and 29 experiments inducing non-visceral (mechanical) pain. Finally, for sample composition-related analyses, we meta-analyzed 22 all-female experiments and 30 all-male experiments (**Figure 2; Table 1**).

**Figure 2.**
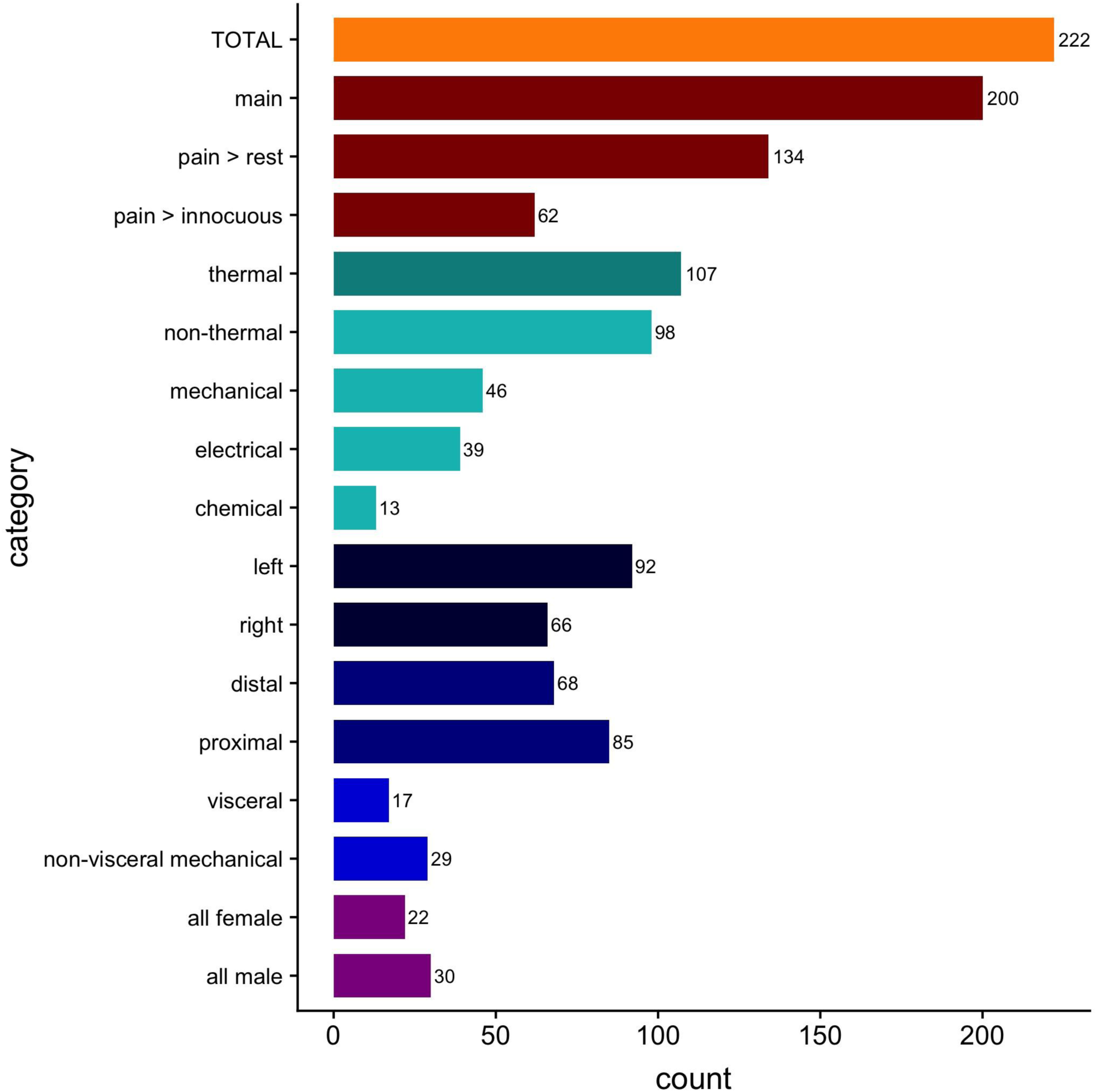
Number of experiments included in each analysis of interest.

**Table 1.**
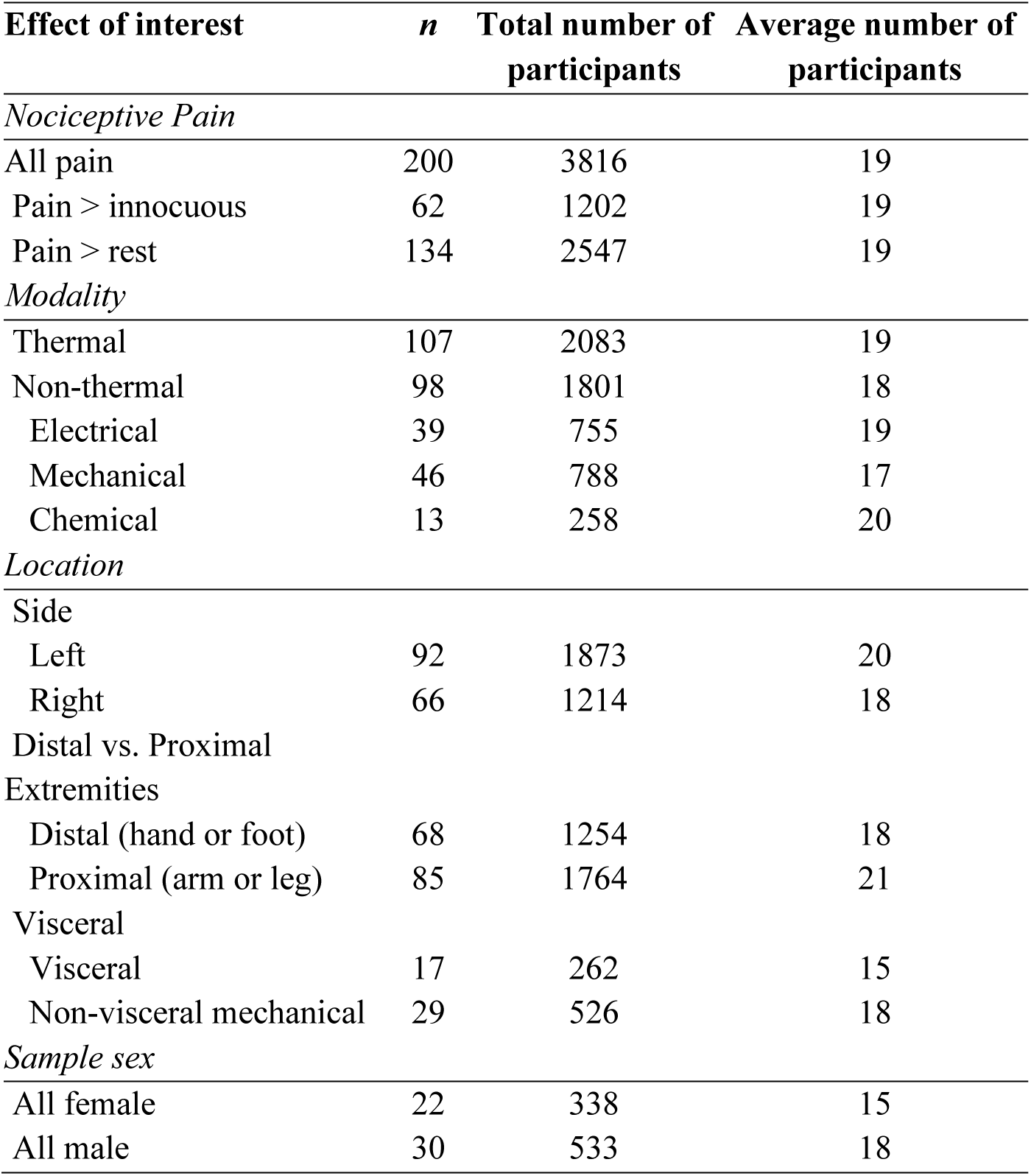
Number of experiments included and number of participants for each effect of interest.

| **Effect of interest** | ***n*** | **Total number of participants** | **Average number of participants** |
| --- | --- | --- | --- |
| *Nociceptive Pain* |  |  |  |
| All pain | 200 | 3816 | 19 |
| Pain > innocuous | 62 | 1202 | 19 |
| Pain > rest | 134 | 2547 | 19 |
| *Modality* |  |  |  |
| Thermal | 107 | 2083 | 19 |
| Non-thermal | 98 | 1801 | 18 |
| Electrical | 39 | 755 | 19 |
| Mechanical | 46 | 788 | 17 |
| Chemical | 13 | 258 | 20 |
| *Location* |  |  |  |
| Side |  |  |  |
| Left | 92 | 1873 | 20 |
| Right | 66 | 1214 | 18 |
| Distal vs. Proximal |  |  |  |
| Extremities |  |  |  |
| Distal (hand or foot) | 68 | 1254 | 18 |
| Proximal (arm or leg) | 85 | 1764 | 21 |
| Visceral |  |  |  |
| Visceral | 17 | 262 | 15 |
| Non-visceral mechanical | 29 | 526 | 18 |
| *Sample sex* |  |  |  |
| All female | 22 | 338 | 15 |
| All male | 30 | 533 | 18 |

### 3.1. Main effect of stimuli inducing a sensation of pain

Meta-analysis of pain experiments from all studies (*n* = 200), which included experiments using the contrasts “pain > rest” and “pain > innocuous” as well as parametric modulation of pain, revealed significant pain-related convergence of activation in four large clusters, with peak activation magnitudes located in the right supramarginal gyrus (inferior parietal lobule, IPL), right midcingulate cortex (MCC), right precentral gyrus, and left cerebellum (**Figure 3; Table 2**). Further examination of these large clusters revealed activation in the bilateral thalamus, bilateral SMA, bilateral pre-SMA, bilateral putamen, bilateral caudate, bilateral brainstem, bilateral amygdala, left supramarginal gyrus/IPL (insula) and left MCC. Post-hoc analysis confirmed these results were not significantly impacted by any individual experiment (see **Supplementary Table 1**).

**Figure 3.**
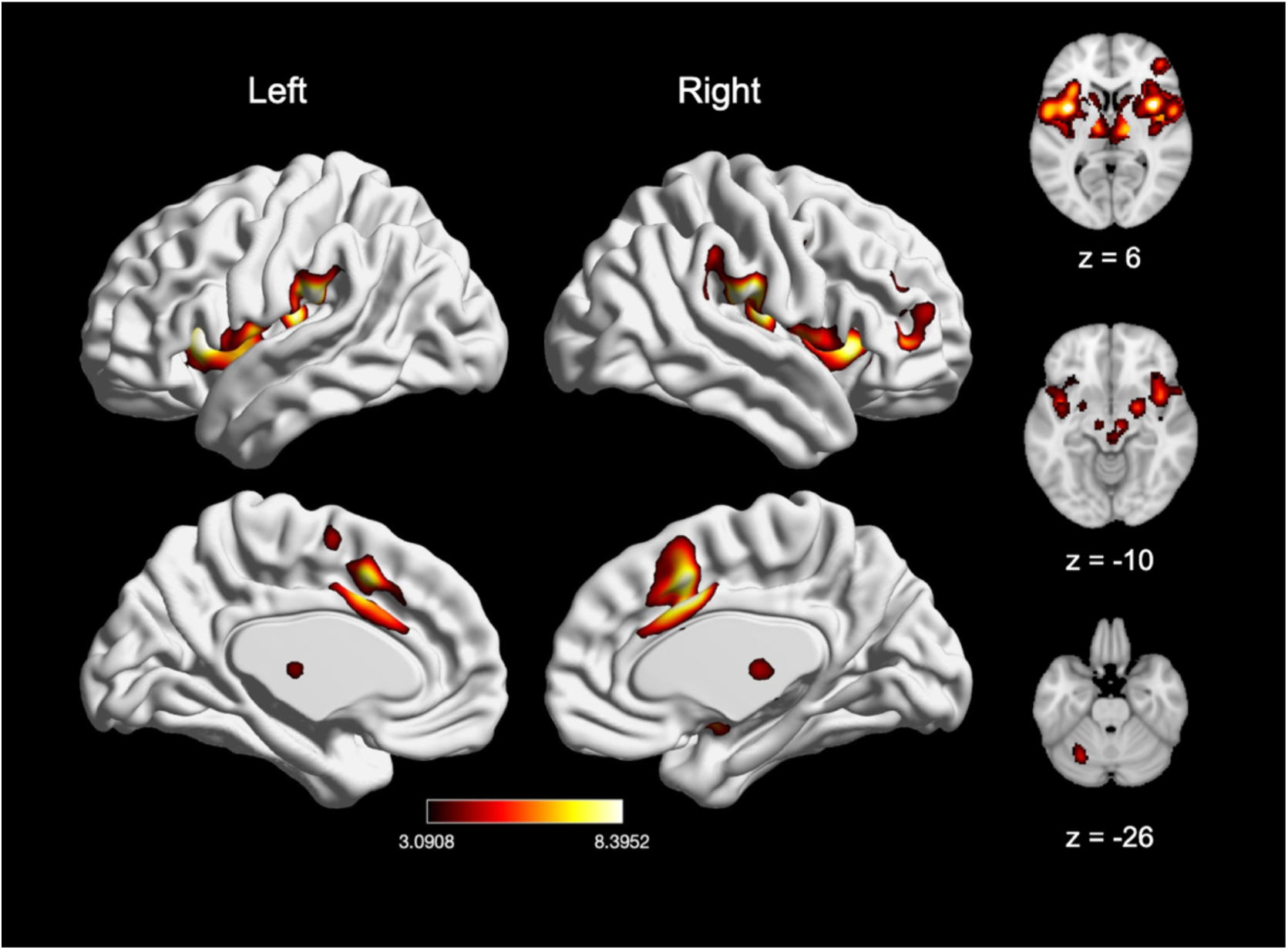
Main effect of experimental induction of acute pain. (A) Main effect of experiencing nociceptive pain across all 200 experiments (includes studies with the contrasts “pain > rest” and “pain > innocuous”, as well as studies examining parametric modulation of pain. (B) Main effect of experiments with “pain > innocuous” contrast (*n* = 62). Coordinates and statistics for significant clusters are shown in **Table 2**.

**Table 2.** Peaks of convergence of activation for meta-analyses related to acute nociceptive pain.

| Effect of Interest | Voxels | X | Y | Z | Macroanatomical<br>Label | Cytoarchitectonic<br>Label | Z-score |
| --- | --- | --- | --- | --- | --- | --- | --- |
| Main effect: |  |  |  |  |  |  |  |
| Pain |  |  |  |  |  |  |  |
|  | 14522 | 58 | -24 | 22 | R Supramarginal Gyrus | Area PFop (IPL) | 8.52 |
|  | 3181 | 6 | 12 | 38 | R MCC |  | 8.48 |
|  | 325 | -32 | -56 | -34 | L Cerebellum | Lobule VI (Hem) | 5.39 |
|  | 238 | 48 | 4 | 42 | R Precentral Gyrus |  | 5.03 |

*Note.* Cluster coordinates are reported in MNI stereotaxic space. Macroanatomical labels used the SPM Anatomy Toolbox and Harvard-Oxford Structural Atlas (MCC = midcingulate cortex). Cytoarchitectonic labels were made using the SPM Anatomy Toolbox (IPL = inferior parietal lobe).

We next evaluated experiments reporting a “pain > rest” contrast (*n =* 134) and “pain > innocuous” contrast (*n* = 62), as “pain > innocuous” contrasts controlled for the impact of sensorimotor stimulation (e.g., touch). In experiments reporting a “pain > rest” contrast, we found significant activation in seven clusters, with peak activation magnitudes located in bilateral IPL (right supramarginal gyrus and left intraparietal sulcus), bilateral pre-SMA, right MCC, right precentral gyrus, left insula, left thalamus, and left cerebellum (**Figure 4A; Table 3**). Activation in these clusters also comprised of bilateral SMA, bilateral pre-SMA, bilateral putamen, bilateral caudate, right brainstem, right insula, right middle frontal gyrus, right amygdala, left MCC, and left precentral gyrus. In experiments using a “pain > innocuous” contrast, we found significant activation in four clusters, with peak activation magnitudes located in bilateral insula, right MCC, and right thalamus (**Figure 4B; Table 3**). These clusters were also comprised of the bilateral putamen, bilateral SMA, left MCC, left thalamus, right amygdala, right precentral gyrus and right pre-SMA. Due to statistically significant differences in number of experiments reporting these contrasts (χ^2^(1) = 27.33, *p* < 0.001), we did not perform an explicit contrast between experiments reporting a “pain > rest” contrast and a “pain > innocuous” contrast.

**Figure 4.**
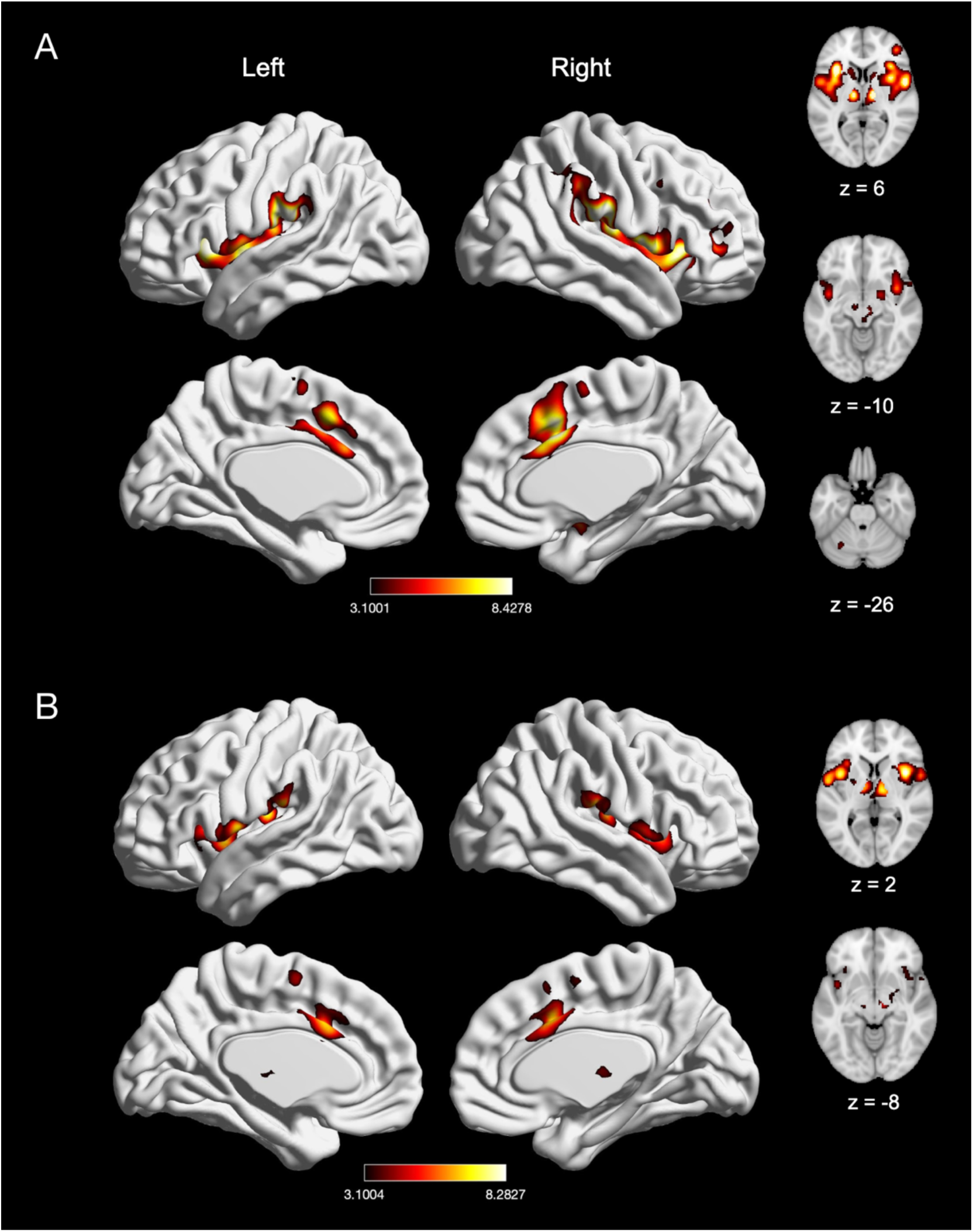
Meta-analytic effect of experiments using “pain > rest” and “pain > innocuous” contrast. (A) Main effect of experiments using “pain > rest” contrast (*n* = 134). (B) Main effect of experiments with “pain > innocuous” contrast (*n* = 62). Coordinates and statistics for significant clusters are shown in **Table 3**.

**Table 3.** Peaks of convergence of activation for meta-analyses related to acute nociceptive pain experiments using “pain > rest” contrast and “pain > innocuous” contrast.

| Effect of Interest | Voxels | X | Y | Z | Macroanatomical<br>Label | Cytoarchitectonic<br>Label | Z-score |
| --- | --- | --- | --- | --- | --- | --- | --- |
| Main effect: |  |  |  |  |  |  |  |
| Pain > rest |  |  |  |  |  |  |  |
|  | 6936 | 58 | -24 | 22 | R Supramarginal Gyrus | Area PFop (IPL) | 8.43 |
|  | 4641 | -34 | 18 | 4 | L Insula Lobe |  | 8.39 |
|  | 2556 | 6 | 12 | 38 | R MCC |  | 8.37 |
|  | 439 | -10 | -14 | 2 | L Thalamus | Thal: Prefrontal | 8.35 |
|  | 164 | -46 | -40 | 44 | L Inferior Parietal<br>Lobule | Area hIP2 (IPS) | 4.71 |
|  | 135 | -30 | -56 | -32 | L Cerebellum (VI) | Lobule VI (Hem) | 4.57 |
|  | 134 | 48 | 4 | 38 | R Precentral Gyrus |  | 4.59 |
| Main effect: |  |  |  |  |  |  |  |
| Pain > innocuous |  |  |  |  |  |  |  |
|  | 2301 | 38 | 8 | 6 | R Insula Lobe |  | 8.28 |
|  | 2287 | -38 | -18 | 14 | L Insula Lobe | Area Ig2 | 7.46 |
|  | 1521 | 6 | 10 | 38 | R MCC |  | 7.58 |
|  | 1005 | 10 | -12 | 4 | R Thalamus | Thal: Prefrontal | 6.88 |

*Note.* Cluster coordinates are reported in MNI stereotaxic space. Macroanatomical labels used the SPM Anatomy Toolbox and Harvard-Oxford Structural Atlas (MCC = midcingulate cortex). Cytoarchitectonic labels were made using the SPM Anatomy Toolbox (IPL = inferior parietal lobe; Thal = thalamus).

### 3.2. Effect of stimulus modality

#### 3.2.1. Thermal and non-thermal pain

To examine modality-specific effects of pain induction, we first compared experiments that induced thermal pain (*n* = 107) with experiments that induced non-thermal pain (*n =* 98). Experiments inducing thermal pain showed convergence of activation in five clusters, with peak activation magnitudes located in right Rolandic operculum/IPL (posterior insula), right MCC, right middle frontal gyrus, right precentral gyrus, and left cerebellum (**Figure 5A; Table 4**). Further examination of these large clusters revealed activation in bilateral putamen, bilateral caudate, bilateral brainstem, bilateral SMA, bilateral pre-SMA, left Rolandic operculum/IPL, left MCC, and right amygdala. Experiments inducing non-thermal pain also showed activation in five clusters, with peak activation magnitudes located in right Rolandic operculum, right MCC, right thalamus, right middle frontal gyrus, and left postcentral gyrus (SII) (**Figure 5B; Table 4**). Further examination of these large clusters revealed activation in bilateral SMA, bilateral putamen, bilateral pre-SMA, left MCC, and left thalamus. An explicit contrast between thermal and non-thermal pain experiments revealed significantly stronger convergence of activation in bilateral MCC for thermal experiments, and stronger convergence in the right insula and left Rolandic operculum in non-thermal experiments (**Table 4**). Conjunction analyses revealed widespread overlap between thermal and non-thermal pain experiments in six large clusters, with peak activation magnitudes located in bilateral supramarginal gyrus (SII), bilateral thalamus, right MCC, and right middle frontal gyrus (**Table 4**). These large clusters also included the bilateral putamen, bilateral SMA, left MCC, and right amygdala.

**Figure 5.**
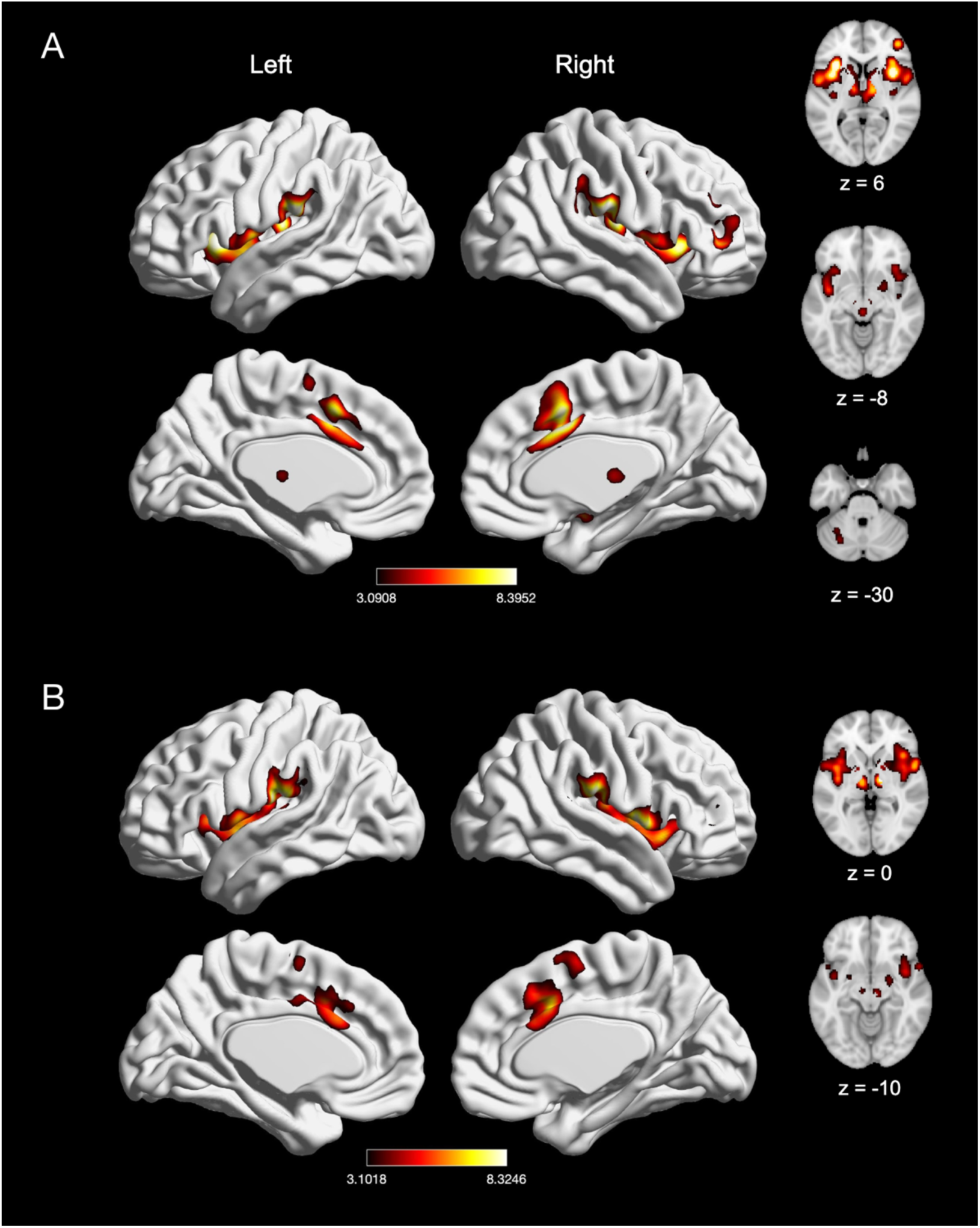
The meta-analytic effects of thermal and non-thermal nociceptive pain induction. (A) Main effect of thermal pain experiments (*n* = 107). (B) Main effect of non-thermal pain experiments (*n* = 98). Coordinates and statistics for significant clusters associated with the main effect of thermal and non-thermal pain (as well as the between-experiment contrast and conjunction of thermal and non-thermal pain experiments) are shown in **Table 4**.

**Table 4.**
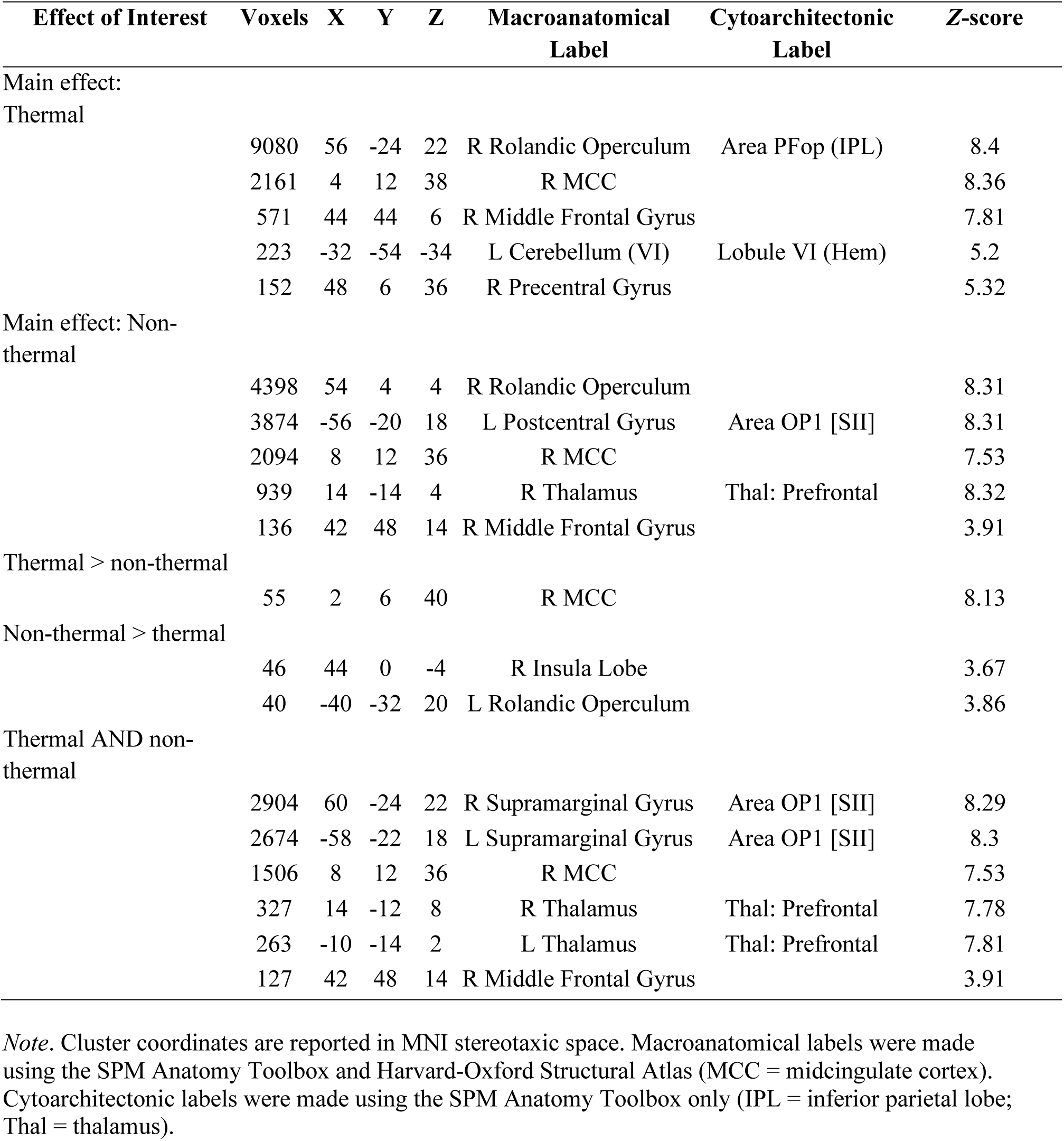
Peaks of convergence of activation for meta-analyses related to acute thermal and non-thermal nociceptive pain.

#### 3.2.2. Electrical, mechanical, and chemical pain

We next examined the effect of electrical (*n =* 39), mechanical (*n =* 46), and chemical (*n* = 13) pain induction (**Figure 6** and **Figure 7**). Experiments inducing electrical stimulation to evoke pain showed convergence of activation in five large clusters, with peak activation magnitudes located in bilateral thalamus, right MCC, right Rolandic operculum, and left postcentral gyrus (SII) (**Figure 6A; Table 5**). Experiments inducing mechanical pain showed convergence of activation in seven large clusters, with peak activation magnitudes located in bilateral insula, bilateral supramarginal gyrus (consisting of SII and IPL), bilateral thalamus, and right MCC (**Figure 6B; Table 5**). Further examination of these large clusters associated with mechanical pain revealed activation including the bilateral putamen and left MCC. An explicit contrast between electrical and mechanical pain experiments revealed greater activation in the right Rolandic operculum, right thalamus, and right superior temporal gyrus in electrical pain experiments. In no cases was greater activation seen for mechanical pain experiments (**Table 5**). Conjunction analyses revealed widespread overlap of convergence of activation between electrical and mechanical pain experiments in seven large clusters, with peak activation magnitudes located in bilateral insula, bilateral SII (left postcentral gyrus and right supramarginal gyrus), bilateral thalamus, and right MCC (**Table 5**).

**Figure 6.**
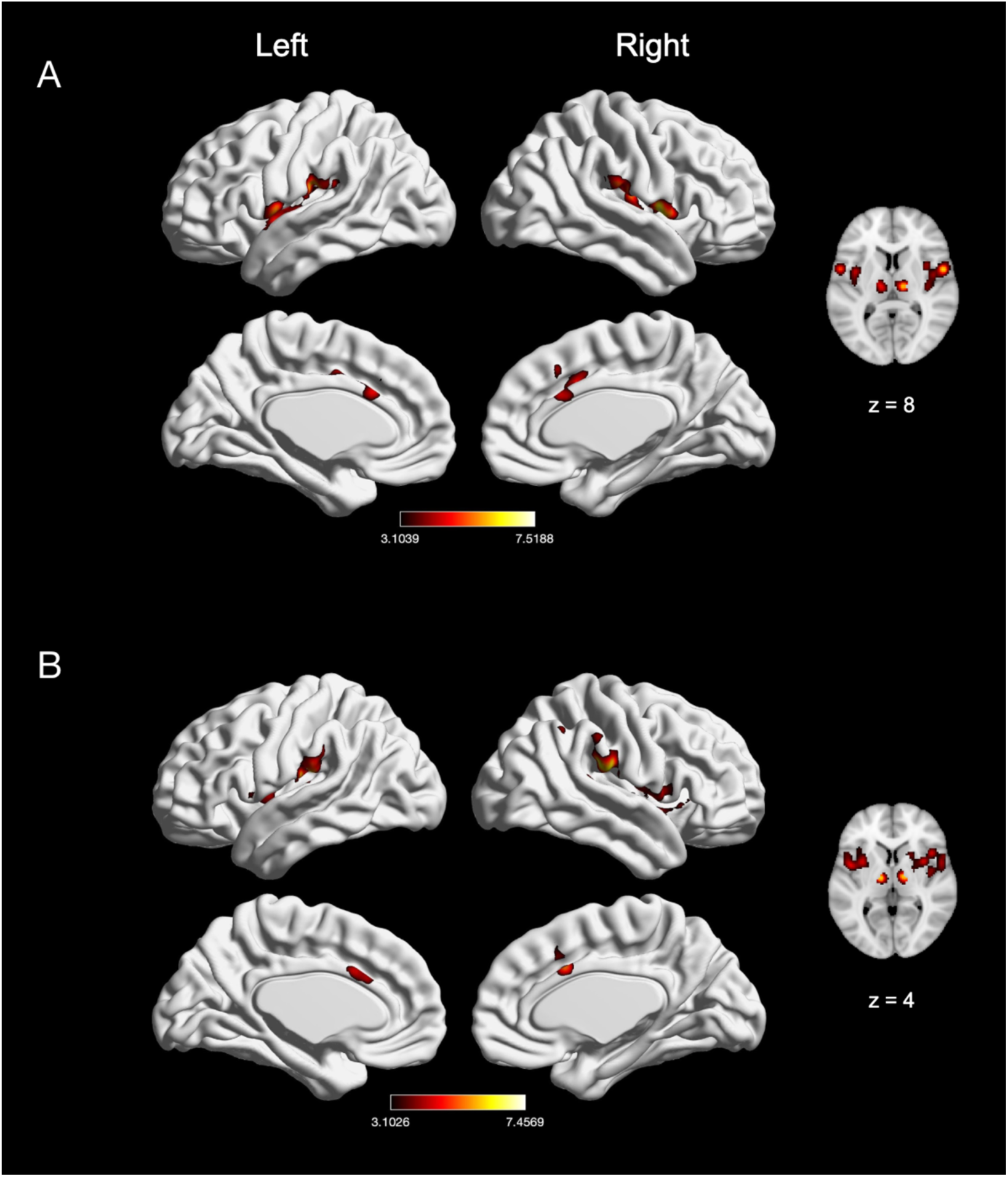
Effects of induction of electrically-evoked and mechanical nociceptive pain. (A) Main effect of electrically-evoked pain experiments (*n* = 39). (B) Main effect of mechanical pain experiments (*n* = 46). Coordinates and statistics for significant clusters associated with the main effect of electrical and mechanical pain (as well as the between-experiment contrast and conjunction of electrical and mechanical pain experiments) are shown in **Table 5**.

**Figure 7.**
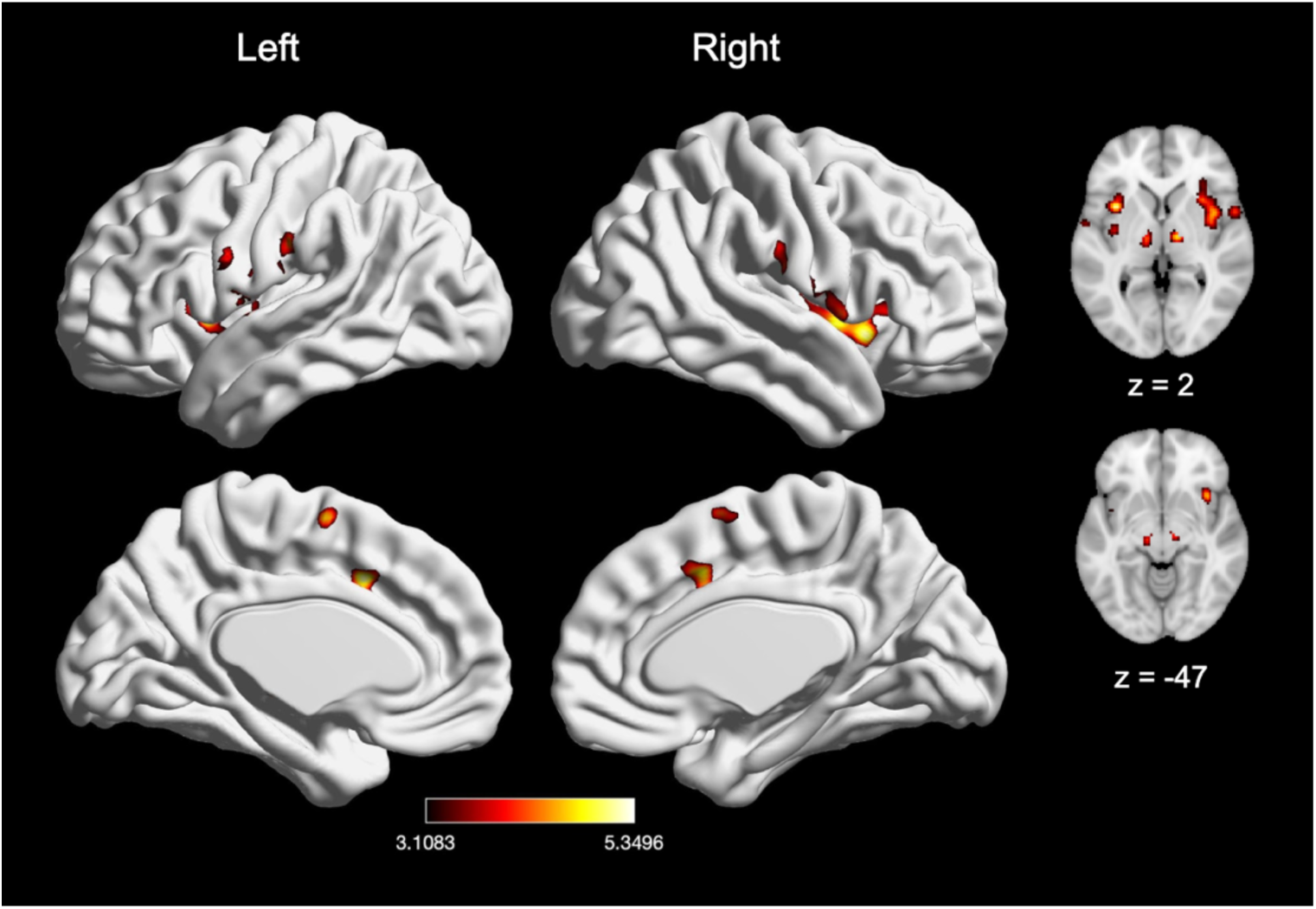
Main effect of meta-analysis of nociceptive chemical pain experiments (*n* = 13). Coordinates and statistics for significant clusters associated with the main effect of chemical pain are shown in Table 5.

**Table 5.** Peaks of convergence of activation for meta-analyses related to acute electrical and mechanical nociceptive pain.

| Effect of Interest | Voxels | X | Y | Z | Macroanatomical Label | Cytoarchitectonic Label | Z-score |
| --- | --- | --- | --- | --- | --- | --- | --- |
| Main effect: |  |  |  |  |  |  |  |
| Electrical |  |  |  |  |  |  |  |
|  | 1901 | 54 | 2 | 6 | R Rolandic Operculum |  | 7.52 |
|  | 1514 | -56 | -20 | 20 | L Postcentral Gyrus | Area OP1 [SII] | 6.17 |
|  | 729 | 10 | 12 | 38 | R MCC |  | 4.87 |
|  | 249 | 14 | -16 | 6 | R Thalamus | Thal: Prefrontal | 6.59 |
|  | 202 | -12 | -16 | 8 | L Thalamus | Thal: Prefrontal | 5.27 |
| Main effect: |  |  |  |  |  |  |  |
| mechanical |  |  |  |  |  |  |  |
|  | 1354 | 34 | 6 | 8 | R Insula Lobe |  | 5.63 |
|  | 848 | -44 | 0 | 6 | L Insula Lobe |  | 5.19 |
|  | 814 | -54 | -22 | 16 | L Supramarginal Gyrus | Area OP1 [SII] | 7.39 |
|  | 755 | 58 | -26 | 24 | R Supramarginal Gyrus | Area PFop (IPL) | 7.46 |
|  | 501 | 4 | 14 | 34 | R MCC |  | 5.7 |
|  | 227 | -12 | -14 | 4 | L Thalamus | Thal: Prefrontal | 6.76 |
|  | 164 | 12 | -12 | 2 | R Thalamus | Thal: Prefrontal | 6.9 |
| Electrical > mechanical |  |  |  |  |  |  |  |
|  | 33 | 44 | -20 | 22 | R Rolandic Operculum |  | 3.83 |
|  | 26 | 6 | -18 | 10 | R Thalamus | Thal: Temporal | 3.54 |
|  | 25 | 46 | 2 | -16 | R Superior Temporal Gyrus |  | 3.45 |
| Mechanical > electrical |  |  |  |  |  |  |  |
| *No suprathreshold clusters* |  |  |  |  |  |  |  |
| Electrical AND mechanical |  |  |  |  |  |  |  |
|  | 466 | 36 | 6 | 10 | R Insula Lobe |  | 4.76 |
|  | 462 | -56 | -20 | 18 | L Postcentral Gyrus | Area OP1 [SII] | 6.06 |
|  | 334 | 60 | -22 | 20 | R Supramarginal Gyrus | Area OP1 [SII] | 5.47 |
|  | 275 | -36 | 4 | 12 | L Insula Lobe |  | 5.05 |
|  | 243 | 8 | 14 | 36 | R MCC |  | 4.62 |
|  | 113 | -14 | -16 | 8 | L Thalamus | Thal: Prefrontal | 5.01 |
|  | 98 | 14 | -14 | 6 | R Thalamus | Thal: Prefrontal | 5.8 |

*Note.* Cluster coordinates are reported in MNI stereotaxic space. Macroanatomical labels were made using the SPM Anatomy Toolbox and Harvard-Oxford Structural Atlas (MCC = midcingulate cortex). Cytoarchitectonic labels were made using the SPM Anatomy Toolbox only (Thal = thalamus).

Finally, chemical pain experiments showed convergence of activation in ten clusters, with peak activation magnitudes located in ventral aspects of the brainstem, bilateral insula (with two clusters in the left insula), bilateral thalamus, left Rolandic operculum, left MCC, left SMA, and right postcentral gyrus/IPL (**Figure 7; Table 6**). Further examination of these large clusters revealed activation within the right MCC and midbrain. There were not a sufficient number of chemical pain induction comparisons to contrast chemical pain-related activation with electrical stimulation-provoked pain (*X^2^*(1) = 13*, p* < 0.001), mechanical pain (*X^2^*(1) = 18.45*, p* < 0.001), or thermal pain (*X^2^*(1) = 73.63*, p* < 0.001).

**Table 6.**
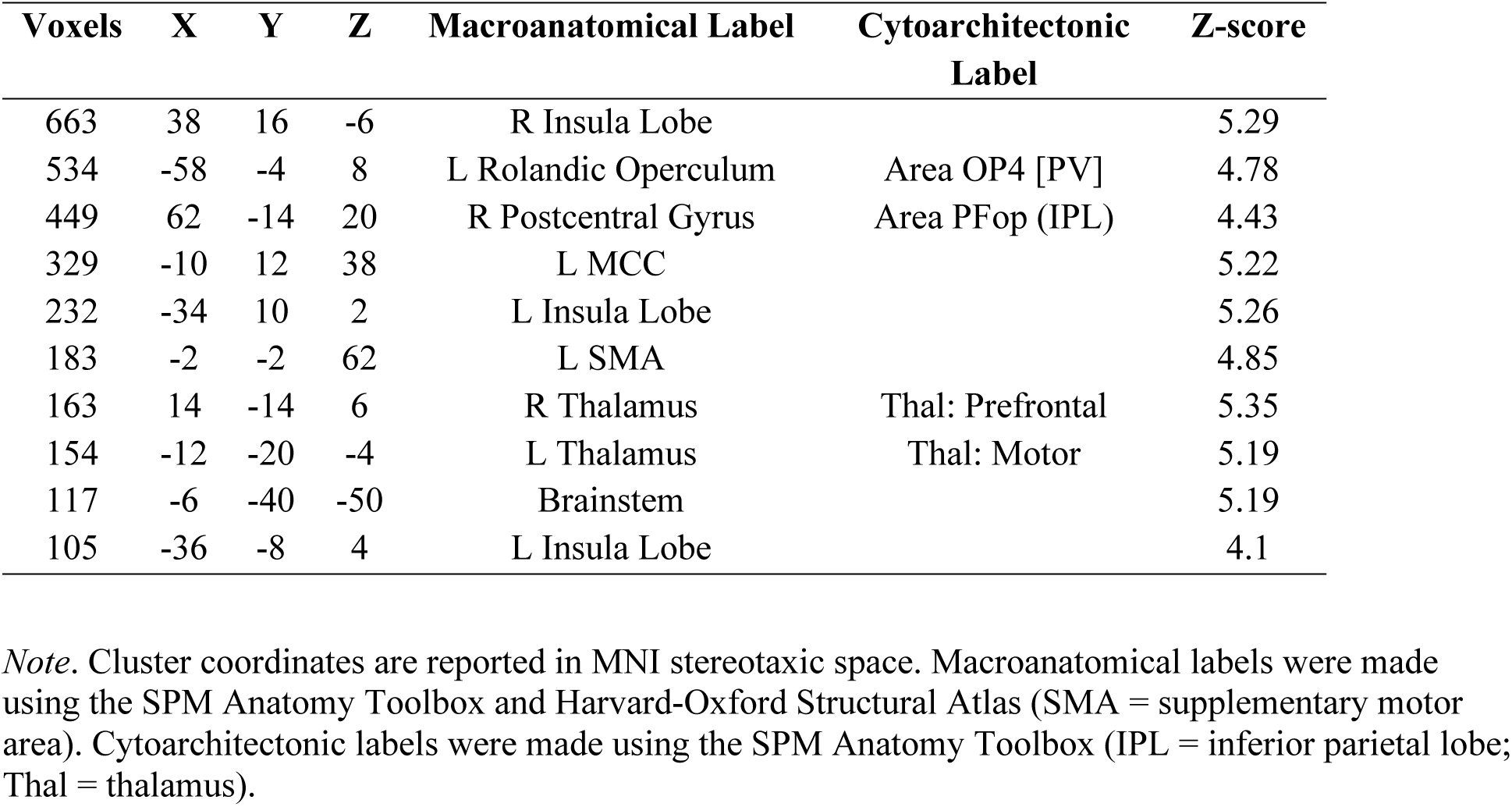
Peaks of convergence of activation for meta-analyses related to acute nociceptive chemical pain

### 3.3. Effect of stimulus location

#### 3.3.1. Laterality

To examine location-specific effects of nociceptive pain induction, we compared experiments that induced left-sided pain (*n* = 92) with experiments that induced right-sided pain (*n* = 66). Experiments inducing left-sided pain showed convergence of activation in six clusters, with peak activation magnitudes located in right Rolandic operculum, right MCC, right middle frontal gyrus, and right postcentral gyrus, left supramarginal gyrus, and left cerebellum (**Figure 8A; Table 7**). Further examination of these large clusters revealed activation in bilateral insula, bilateral thalamus, bilateral pre-SMA, left MCC, right amygdala, right pallidum and the brainstem. Experiments inducing right-sided pain showed convergence of activation in nine clusters, with peak activation magnitudes located in bilateral thalamus, left Rolandic operculum, right supramarginal gyrus, right MCC, right middle frontal gyrus, right IPL, right precentral gyrus and right insula (**Figure 8B, Table 7**). Further examination of these large clusters revealed activation in bilateral putamen, bilateral SMA, bilateral pre-SMA, left MCC, left insula, and right precentral gyrus.

**Figure 8.**
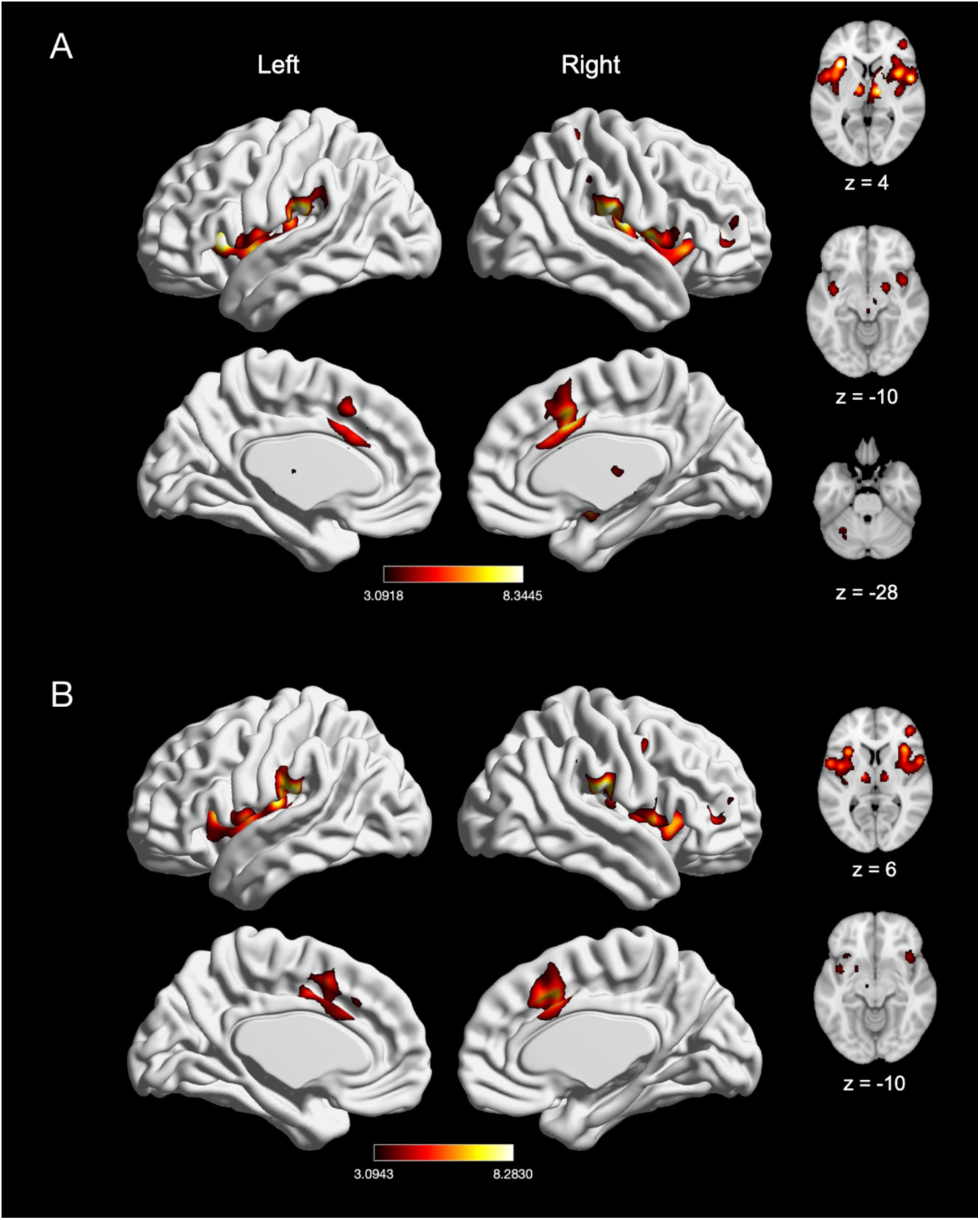
The meta-analytic effects of left-sided and right-sided pain induction. (A) Main effect of left-sided pain experiments (*n* = 92). (B) Main effect of right-sided pain experiments (*n* = 66). Coordinates and statistics for significant clusters associated with the main effect of left-sided and right-sided pain (as well as the between-experiment contrast and conjunction of left-sided and right-sided pain experiments) are shown in **Table7**.

**Table 7.** Peaks of convergence of activation for meta-analyses related to left-sided and right-sided acute nociceptive pain.

| Effect of Interest | Voxels | X | Y | Z | Macroanatomical Label | Cytoarchitectonic Label | Z-score |
| --- | --- | --- | --- | --- | --- | --- | --- |
| Main effect: |  |  |  |  |  |  |  |
| Left |  |  |  |  |  |  |  |
|  | 5034 | 40 | -16 | 16 | R Rolandic Operculum | Area OP3 [VS] | 8.34 |
|  | 3096 | -60 | -24 | 20 | L Supramarginal Gyrus | Area PFop (IPL) | 8.31 |
|  | 1445 | 6 | 12 | 38 | R MCC |  | 8.29 |
|  | 236 | 44 | 44 | 6 | R Middle Frontal Gyrus |  | 5.73 |
|  | 137 | -32 | -56 | -32 | L Cerebellum (VI) | Lobule VI (Hem) | 4.62 |
|  | 125 | 24 | -44 | 68 | R Postcentral Gyrus | Area 1 | 5.87 |
| Main effect: |  |  |  |  |  |  |  |
| Right |  |  |  |  |  |  |  |
|  | 2944 | -40 | -18 | 14 | L Rolandic Operculum | Area Ig2 | 8.27 |
|  | 1743 | 38 | 20 | 4 | R Insula Lobe |  | 7.61 |
|  | 1737 | 6 | 12 | 38 | R MCC |  | 7.63 |
|  | 689 | 58 | -22 | 22 | R Supramarginal Gyrus | Area PFop (IPL) | 8.28 |
|  | 301 | 44 | 44 | 6 | R Middle Frontal Gyrus |  | 5.99 |
|  | 224 | -12 | -14 | 4 | L Thalamus | Thal: Prefrontal | 6.16 |
|  | 217 | 50 | 2 | 44 | R Precentral Gyrus |  | 5.31 |
|  | 170 | 14 | -14 | 6 | R Thalamus | Thal: Prefrontal | 5.06 |
|  | 143 | 52 | -38 | 46 | R Inferior Parietal Lobule | Area PFm (IPL) | 4.1 |
| Left > right |  |  |  |  |  |  |  |
| *No suprathreshold clusters* |  |  |  |  |  |  |  |
| Right > left |  |  |  |  |  |  |  |
|  | 45 | -44 | -22 | 16 | L Rolandic Operculum | Area OP1 [SII] | 6.8 |
| Left AND right |  |  |  |  |  |  |  |
|  | 1437 | -32 | 18 | 6 | L Insula Lobe |  | 7.11 |
|  | 1318 | 38 | 20 | 0 | R Insula Lobe |  | 6.76 |
|  | 1116 | 6 | 12 | 38 | R MCC |  | 7.63 |
|  | 608 | 58 | -22 | 22 | R Supramarginal Gyrus | Area PFop (IPL) | 8.28 |
|  | 542 | -58 | -26 | 20 | L Supramarginal Gyrus | Area PFop (IPL) | 8.08 |
|  | 178 | 44 | 44 | 6 | R Middle Frontal Gyrus |  | 5.73 |
|  | 167 | 14 | -14 | 6 | R Thalamus | Thal: Prefrontal | 5.06 |
|  | 158 | -12 | -14 | 4 | L Thalamus | Thal: Prefrontal | 6.16 |

*Note.* Cluster coordinates are reported in MNI stereotaxic space. Macroanatomical labels were made using the SPM Anatomy Toolbox and Harvard-Oxford Structural Atlas (MCC = midcingulate cortex). Cytoarchitectonic labels were made using the SPM Anatomy Toolbox only (IPL = inferior parietal lobe; Thal = thalamus; VS = ventral somatosensory).

An explicit contrast between left- and right-sided stimulation experiments did not reveal any clusters with stronger convergence of activation in left-sided pain induction experiments but did reveal stronger convergence of activation in left Rolandic operculum (SII) in right-sided pain induction experiments (**Table 7**). Given that we did not see involvement of SI in right-sided pain induction based on these analyses (despite reports of SI involvement in pain (Bushnell, 1999)), we also separately examined unthresholded maps and found involvement of right SI, suggesting that while SI did not appear prominently in our explicit contrast, there may still be some laterality in SI that is harder to detect. SI was particularly prominent when the stimulation site was more tightly aligned across studies (e.g., left-sided arm stimulation).

Finally, conjunction analyses revealed widespread overlap of convergence of activation between left-sided and right-sided experiments in eight clusters, with peak activation magnitudes located in bilateral insula, bilateral supramarginal gyrus (consisting of IPL), bilateral thalamus, right MCC, and right middle frontal gyrus (**Table 7**). Further evaluation of these clusters revealed activation in the left MCC, right putamen, and right SMA.

#### 3.3.2. Distal and proximal extremities

We next compared experiments inducing distal nociceptive pain in the hand or foot (*n* = 68) with experiments inducing proximal pain in the arm or leg (*n* = 85). Experiments inducing distal pain in the hand or foot also showed convergence of activation in six clusters, with peak activation magnitudes located in bilateral IPL (consisting of right Rolandic operculum and left supramarginal gyrus), bilateral thalamus, right MCC, and right middle frontal gyrus (**Figure 9A; Table 8**). Further examination of these large clusters revealed activation in bilateral insula, bilateral amygdala, bilateral pre-SMA, and right SMA. Experiments inducing proximal pain showed convergence of activation in six clusters, with peak activation magnitudes located in right Rolandic operculum, right MCC, right middle frontal gyrus, right precentral gyrus, right post central gyrus, and left insula (**Figure 9B; Table 8**). These large clusters included the bilateral pre-SMA, bilateral subnuclei of the striatum, right insula, right SMA, and left MCC. An explicit contrast between proximal and distal stimulation experiments did not reveal differential patterns of activation, while conjunction analyses revealed widespread overlap convergence of activation in eight clusters (**Table 8**). These eight clusters included bilateral thalamus, bilateral IPL (represented by left supramarginal gyrus and right operculum), right MCC (with spread to left MCC), right middle frontal gyrus, right pallidum, and left insula.

**Figure 9.**
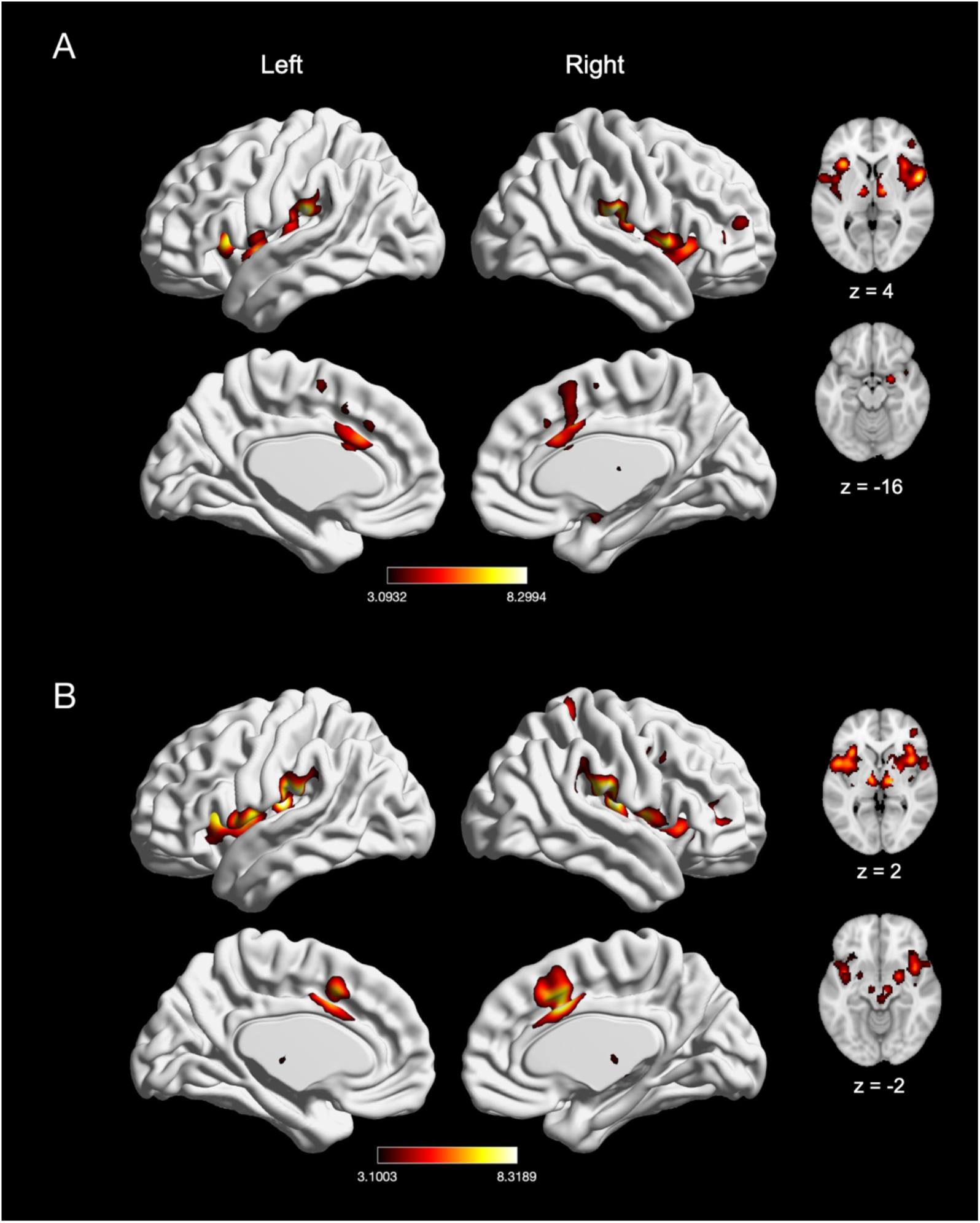
The meta-analytic effects of distal and proximal nociceptive pain. (A) Main effect of experiments inducing acute nociceptive pain in the distal extremities (*n* = 68). (B) Main effect of experiments inducing pain in the proximal extremities (*n* = 85). Coordinates and statistics for significant clusters associated with the main effect of distal and proximal pain (as well as the between-experiment contrast and conjunction of distal and proximal pain experiments) are shown in **Table 8**.

**Table 8.** Peaks of convergence of activation for meta-analyses related to distal and proximal acute nociceptive pain.

| Effect of Interest | Voxels | X | Y | Z | Macroanatomical Label | Cytoarchitectonic Label | Z-score |
| --- | --- | --- | --- | --- | --- | --- | --- |
| Main effect: |  |  |  |  |  |  |  |
| Distal |  |  |  |  |  |  |  |
|  | 3044 | 56 | -26 | 22 | R Rolandic Operculum | Area PFop (IPL) | 8.3 |
|  | 2579 | -60 | -24 | 20 | L Supramarginal Gyrus | Area PFop (IPL) | 8.28 |
|  | 1384 | 4 | 10 | 38 | R MCC |  | 6.46 |
|  | 605 | 12 | -12 | 4 | R Thalamus | Thal: Prefrontal | 6.68 |
|  | 222 | 46 | 42 | 6 | R Middle Frontal Gyrus |  | 4.72 |
|  | 162 | -10 | -12 | 2 | L Thalamus | Thal: Prefrontal | 6.21 |
| Main effect: |  |  |  |  |  |  |  |
| Proximal |  |  |  |  |  |  |  |
|  | 4319 | 38 | -18 | 16 | R Rolandic Operculum | Area OP2 [PIVC] | 8.32 |
|  | 3261 | -36 | 4 | 8 | L Insula Lobe |  | 8.31 |
|  | 1696 | 6 | 12 | 38 | R MCC |  | 8.31 |
|  | 227 | 44 | 44 | 6 | R Middle Frontal Gyrus |  | 6.05 |
|  | 223 | 48 | 6 | 38 | R Precentral Gyrus |  | 5.62 |
|  | 182 | 22 | -44 | 68 | R Postcentral Gyrus | Area 1 | 6.5 |
| Distal > proximal |  |  |  |  |  |  |  |
| *No suprathreshold clusters* |  |  |  |  |  |  |  |
| Proximal > distal |  |  |  |  |  |  |  |
| *No suprathreshold clusters* |  |  |  |  |  |  |  |
| Distal AND proximal |  |  |  |  |  |  |  |
|  | 2041 | 56 | -24 | 22 | R Rolandic Operculum | Area PFop (IPL) | 8.3 |
|  | 1156 | -34 | 20 | 4 | L Insula Lobe |  | 7.25 |
|  | 917 | 4 | 10 | 38 | R MCC |  | 6.46 |
|  | 854 | -60 | -24 | 20 | L Supramarginal Gyrus | Area PFop (IPL) | 8.28 |
|  | 258 | 12 | -12 | 4 | R Thalamus | Thal: Prefrontal | 6.68 |
|  | 153 | -10 | -12 | 2 | L Thalamus | Thal: Prefrontal | 6.21 |
|  | 97 | 46 | 42 | 6 | R Middle Frontal Gyrus |  | 4.69 |
|  | 77 | 16 | 4 | 0 | R Pallidum |  | 4.01 |

*Note.* Cluster coordinates are reported in MNI stereotaxic space. Macroanatomical labels were made using the SPM Anatomy Toolbox and Harvard-Oxford Structural Atlas (MCC = midcingulate cortex). Cytoarchitectonic labels were made using the SPM Anatomy Toolbox only (IPL = inferior parietal lobe; Thal = thalamus).

#### 3.3.3. Effect of acute visceral and non-visceral mechanical nociceptive pain

We compared experiments inducing visceral pain (e.g., distension in the rectum, esophagus, or stomach; *n* = 17) with experiments inducing non-visceral mechanical pain (*n* = 29). Experiments inducing visceral pain showed convergence of activation in five clusters, with peak activation magnitudes located in right supramarginal gyrus, right Rolandic operculum, left putamen, left thalamus, and right MCC (**Figure 10A; Table 9**). Non-visceral mechanical pain experiments showed convergence of activation in ten clusters, with peak activation magnitudes located in bilateral supramarginal gyrus (SII), bilateral insula (with two clusters in the right insula), bilateral thalamus, right Rolandic operculum, right caudate nucleus, and right MCC (**Figure 10B; Table 9**). An explicit contrast between visceral and non-visceral mechanical pain experiments did not reveal differential activation, while conjunction analyses revealed overlap of convergence in two clusters—the right supramarginal gyrus and the left thalamus (**Table 9**).

**Figure 10.**
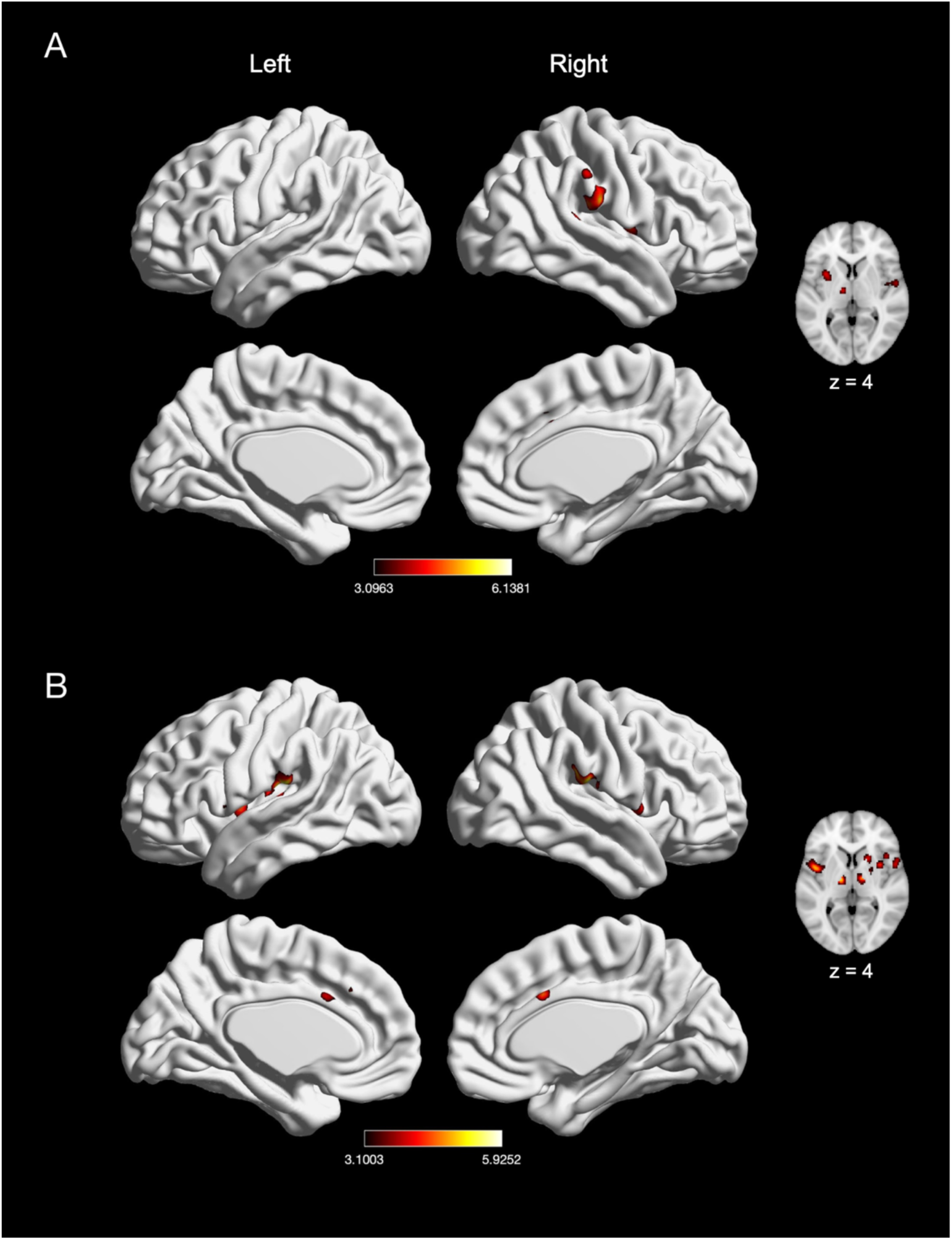
The meta-analytic effects of visceral and non-visceral mechanical pain. (A) Main effect of visceral pain experiments (*n* = 17). (B) Main effect of non-visceral mechanical pain experiments (*n* = 29). Coordinates and statistics for significant clusters associated with the main effect of visceral and non-visceral pain (as well as the between-experiment contrast and conjunction of visceral and non-visceral pain experiments) are shown in **Table 9**.

**Table 9.** Peaks of convergence of activation for meta-analyses related to acute visceral and non-visceral nociceptive pain.

| Effect of Interest | Voxels | X | Y | Z | Macroanatomical Label | Cytoarchitectonic Label | Z-score |
| --- | --- | --- | --- | --- | --- | --- | --- |
| Main effect: |  |  |  |  |  |  |  |
| Visceral |  |  |  |  |  |  |  |
|  | 431 | 60 | -24 | 26 | R Supramarginal Gyrus | Area PFop (IPL) | 6.14 |
|  | 237 | -32 | 0 | -2 | L Putamen |  | 4.76 |
|  | 174 | 54 | -6 | 8 | R Rolandic Operculum | Area OP4 [PV] | 4.66 |
|  | 142 | -8 | -14 | -2 | L Thalamus | Thal: Prefrontal | 4.89 |
|  | 91 | 8 | 16 | 36 | R MCC |  | 4.2 |
| Main effect: |  |  |  |  |  |  |  |
| Non-visceral mechanical |  |  |  |  |  |  |  |
|  | 606 | -56 | -24 | 18 | L Supramarginal Gyrus | Area OP1 [SII] | 5.93 |
|  | 394 | -44 | 2 | 4 | L Insula Lobe |  | 5.24 |
|  | 351 | 60 | -22 | 18 | R Supramarginal Gyrus | Area OP1 [SII] | 5.71 |
|  | 272 | 4 | 14 | 34 | R MCC |  | 4.71 |
|  | 185 | 44 | 18 | 0 | R Insula Lobe |  | 4.59 |
|  | 150 | 54 | 8 | 6 | R Rolandic Operculum | Area 44 | 4.31 |
|  | 140 | 36 | 6 | 6 | R Insula Lobe |  | 4.87 |
|  | 128 | 20 | 14 | 4 | R Caudate |  | 4.38 |
|  | 108 | -14 | -14 | 4 | L Thalamus | Thal: Prefrontal | 5.52 |
|  | 100 | 14 | -12 | 4 | R Thalamus | Thal: Prefrontal | 5 |
| Visceral > Non-visceral mechanical |  |  |  |  |  |  |  |
| *No suprathreshold clusters* |  |  |  |  |  |  |  |
| Non-visceral mechanical > visceral |  |  |  |  |  |  |  |
| *No suprathreshold clusters* |  |  |  |  |  |  |  |
| Visceral AND Non-visceral mechanical |  |  |  |  |  |  |  |
|  | 136 | 58 | -28 | 24 | R Supramarginal Gyrus | Area PFcm (IPL) | 5.18 |
|  | 74 | -10 | -14 | 2 | L Thalamus | Thal: Prefrontal | 4.49 |

*Note.* Cluster coordinates are reported in MNI stereotaxic space. Macroanatomical labels were made using the SPM Anatomy Toolbox and Harvard-Oxford Structural Atlas (MCC = midcingulate cortex). Cytoarchitectonic labels were made using the SPM Anatomy Toolbox only (IPL = inferior parietal lobe; Thal = thalamus).

### 3.4. Effect of sex

We examined sex differences in nociceptive pain responses by comparing experiments with an all-female sample (*n* = 22) to those with an all-male sample (*n* = 30). Experiments with an all-female sample showed convergence of activation in five clusters, with peak activation magnitudes located in bilateral IPL (represented by left superior temporal gyrus and right supramarginal gyrus) and bilateral insula (with two clusters in the left insula; see **Figure 11A and Table 10**). Experiments with an all-male sample showed convergence of activation in eight clusters, with peak activation magnitudes located in bilateral insula, bilateral thalamus, right MCC (spreading into the left hemisphere), right temporal pole (spreading into the precentral gyrus), right middle frontal gyrus and right Rolandic operculum (**Figure 11B; Table 10**). An explicit contrast between males and females did not reveal differential activation (**Table 10**), while conjunction analyses revealed overlap in three clusters—bilateral insula and left supramarginal gyrus/IPL (**Table 10**).

**Figure 11.**
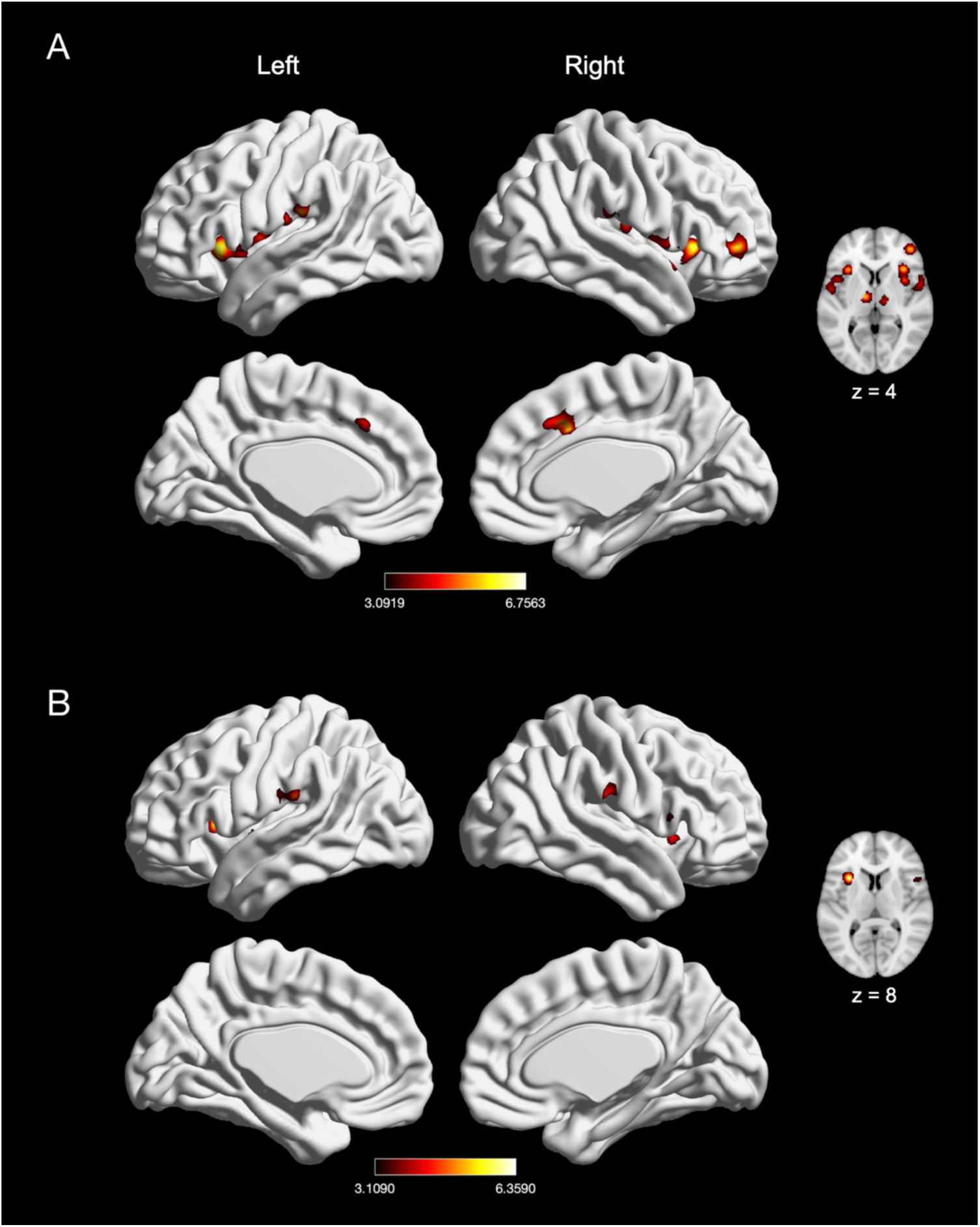
Effect of pain in males and females. (A) Main effect of experiments with an all-female sample (*n* = 22). (B) Main effect of experiments with an all-male sample (*n* = 30). Coordinates and statistics for significant clusters are shown in **Table 10**.

**Table 10.** Peaks of convergence of activation for meta-analyses related to all-female and all-male samples.

| Effect of Interest | Voxels | X | Y | Z | Macroanatomical Label | Cytoarchitectonic Label | Z-score |
| --- | --- | --- | --- | --- | --- | --- | --- |
| Main effect: |  |  |  |  |  |  |  |
| Female |  |  |  |  |  |  |  |
|  | 315 | -32 | 16 | 8 | L Insula Lobe |  | 6.36 |
|  | 249 | -64 | -28 | 22 | L Superior Temporal Gyrus | Area PFop (IPL) | 5.02 |
|  | 213 | 38 | 16 | -2 | R Insula Lobe |  | 4.48 |
|  | 142 | -40 | -4 | 0 | L Insula Lobe |  | 4.27 |
|  | 98 | 62 | -20 | 24 | R Supramarginal Gyrus | Area PFt (IPL) | 4.98 |
| Main effect: |  |  |  |  |  |  |  |
| Male |  |  |  |  |  |  |  |
|  | 1115 | -34 | 20 | 0 | L Insula Lobe |  | 6.76 |
|  | 547 | 8 | 14 | 36 | R MCC |  | 6.39 |
|  | 477 | 36 | 22 | 2 | R Insula Lobe |  | 6.52 |
|  | 374 | 56 | 12 | -4 | R Temporal Pole | Area TE 3 | 5.03 |
|  | 305 | 56 | -26 | 22 | R Rolandic Operculum | Area PFop (IPL) | 4.87 |
|  | 274 | 44 | 46 | 4 | R Middle Frontal Gyrus |  | 6.05 |
|  | 189 | -10 | -12 | 4 | L Thalamus | Thal: Prefrontal | 5.98 |
|  | 169 | 14 | -14 | 8 | R Thalamus | Thal: Prefrontal | 5.59 |
| Female > male |  |  |  |  |  |  |  |
| *No suprathreshold clusters* |  |  |  |  |  |  |  |
| Male > female |  |  |  |  |  |  |  |
| *No suprathreshold clusters* |  |  |  |  |  |  |  |
| Female AND male |  |  |  |  |  |  |  |
|  | 141 | -62 | -26 | 22 | L Supramarginal Gyrus | Area PFop (IPL) | 4.88 |
|  | 88 | -32 | 18 | 4 | L Insula Lobe |  | 5.13 |
|  | 58 | 38 | 18 | -2 | R Insula Lobe |  | 4.14 |

*Note.* Cluster coordinates are reported in MNI stereotaxic space. Macroanatomical labels were made using the SPM Anatomy Toolbox and Harvard-Oxford Structural Atlas (MCC = midcingulate cortex). Cytoarchitectonic labels were made using the SPM Anatomy Toolbox only (IPL = inferior parietal lobe; Thal = thalamus).

## 4. DISCUSSION

We conducted what is to our knowledge the largest fMRI meta-analysis of experimentally induced pain in healthy volunteers to date, applying rigorous type I error control and stringent inclusion criteria reflective of current neuroimaging standard guidelines. We found that painful stimulation by acute noxious/electrical stimuli elicited fMRI activation in a core set of brain regions, irrespective of the specific experimental paradigm. These regions include the thalamus, MCC, SII, insula, as well as portions of the supramarginal gyrus/IPL and Rolandic operculum. In a smaller number of analyses, we observed involvement of the lateral PFC, precentral gyrus, SMA, pre-SMA, cerebellum, the brainstem, basal ganglia (putamen, caudate, and palladium), amygdala, and postcentral gyrus (SI; **Figure 12**).

**Figure 12.**
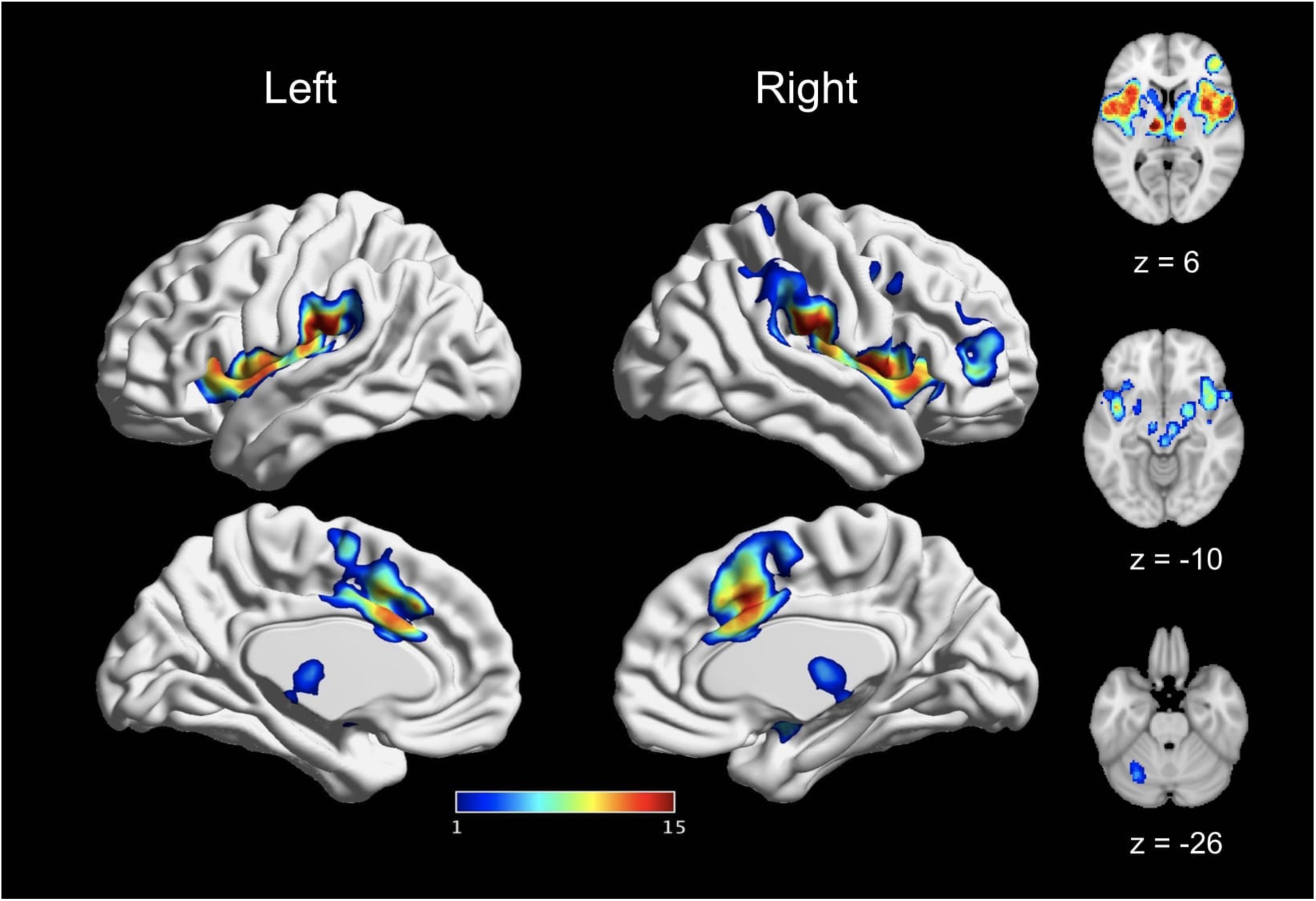
A core set of bran regions recruited by acute pain. This figure depicts the spatial consistency of above-threshold activation convergence cross all reported main effects meta-analyses. The value assigned to each voxel reflects the number of main effects analyses in which it was reported as significant. Values range from 1 (reported significant in only one main effects meta-analyses) to 15 (reported as significant in all main effects meta-analyses). The most consistently activated areas include bilateral thalamus, bilateral insula, bilateral SII, and bilateral MCC.

### 4.1. Core regions activated by pain

The thalamus, SII, MCC, and insula were the most robustly activated brain areas across all experimental paradigms. This finding is consistent with previous meta-analyses (Apkarian et al., 2005; Duerden & Albanese, 2013; Farrell et al., 2005; Friebel et al., 2011; Jensen et al., 2016; Lanz et al., 2011; Peyron et al., 2000; Tanasescu et al., 2016; Tillisch et al., 2011) and with earlier work that delineated the “neurologic pain signature” (Wager et al., 2013). This core set of brain regions has been implicated in sensory-discriminative and affective-motivational aspects of pain (Tracey & Mantyh, 2007; Treede et al., 1999). The thalamus processes and transmits nociceptive information between the spinal cord and cortex (Yen & Lu, 2013; Ab Aziz & Ahmad, 2006) while the SII may reflect higher-order sensory representation, especially information from sensory stimuli requiring more attention (Chen et al., 2008; Ferretti et al., 2003). Here, we found consistent activation of SII even in experiments that include a specific sensorimotor control condition, supporting a role for SII that goes beyond basic sensory processing and concords with previous pain imaging studies demonstrating bilateral SII activation (Mazzola et al., 2006).

The MCC, on the other hand, is strongly implicated in the affective-motivational components of pain, especially in pain response selection (Medford & Critchley, 2010; Shackman et al., 2011; Vogt, 2005, 2016; Vogt, Berger, & Derbyshire, 2003) and has been reported in previous meta-analyses of pain (Farrell et al., 2005; Friebel et al., 2011; Peyron et al., 2000; Tillisch et al., 2011). While other meta-analyses and reviews have highlighted involvement of the anterior cingulate (ACC) in pain (Apkarian et al., 2005; Duerden & Albanese, 2013; Jensen et al., 2016; Lanz et al., 2011; Peyron et al., 2000), it should be noted that our MCC results overlap with what has sometimes been labeled as ACC in these earlier studies. Moreover, the convergence of mid-cingulate activation associated with pain (rather than anterior regions of the cingulate) support delineations of the cingulate cortex with the more mid regions being involved in pain compared to anterior regions (Shackman et al., 2011; Vogt et al., 2016).

The insula is thought to play a more indirect role in pain perception by integrating exteroceptive and interoceptive information into awareness and subjective feelings towards salient information (Craig et al. 2000; Isnard et al., 2011; Craig, 2009; Kurth et al., 2010), especially via its functional connections to the cingulate cortex (Taylor et al., 2009). The posterior portion of the insula in particular has been implicated in processing bodily information (e.g., painful sensations, somatosensory stimulation, interoception) while the anterior insula may be more involved in targeted awareness of salient information (Craig, 2009; Kurth et al., 2010; Menon & Uddin, 2010). As such, our meta-analytic findings of widespread insula activation associated with pain are consistent with involvement of both anterior and posterior insula involved in acute nociceptive pain (Kurth et al., 2010).

### 4.2. Less consistent regions associated with acute pain

In addition to the core set of “pain regions” described above, a broader set of pain-associated regions were somewhat less consistently recruited, emerging only in specific cases. These regions include the lateral PFC, M1, SMA (and portions of the pre-SMA), cerebellum, brainstem, SI, basal ganglia (putamen, caudate, and pallidum), and the amygdala. These areas have been previously reported to have less consistent and possibly more nuanced involvement in pain perception (Apkarian et al., 2005; Duerden & Albanese, 2013; Farrell et al., 2005; Friebel et al., 2011; Lanz et al., 2011; Peyron et al., 2000; Tillisch et al., 2011). Lateral PFC is typically associated with executive control and attention (Bingel & Tracey, 2008; Lorenz et al., 2003; Wiech et al., 2008), while MI, SMA, and cerebellum have typically been associated with execution of motor responses to avoid pain (Apkarian et al., 2005; Peyron et al., 2000; Moulton et al., 2010). The brainstem receives nociceptive input from the spinal cord and trigeminal nucleus and is also involved in the descending modulation of pain (Basbaum & Fields, 1978; Stamford, 1995) while basal ganglia involvement in pain may reflect modulation of multisensory integration related to pain (Borsook et al., 2010; Chudler & Dong, 1995). We observed convergent amygdala activation in four main effects analyses; the amygdala receives dense projections from nociresponsive neurons in the lateral parabrachial nucleus (Jasmin et al., 1997), has been associated with affective modulation of pain and salience detection, and has been reported in both experimental pain and chronic pain experiments (Simons et al., 2014; Borsook et al., 2013; Fernando et al., 2013).

Finally, we did not find consistent involvement of SI in the experimentally-induced pain despite its prior inclusion in a putative “pain matrix” (Iannetti & Mouraux, 2010; Legrain et al., 2011). However, we were able to detect SI involvement in pain in our meta-analyses of left-sided pain and of pain induced in proximal extremities. This less consistent involvement of SI is not surprising in the context of previous reviews have similarly reported less consistent activation of SI in response to painful stimulation compared to other areas (Bushnell et al., 1999; Friebel et al., 2011; Lanz et al., 2011; Peyron et al., 2000). Though SI is known to be critical for the the sensory-discriminative aspect of pain (Bushnell et al., 1999; Duerden & Albanese, 2013; Friebel et al., 2011; Peyron et al., 2000), this inconsistency of SI’s specific involvement in pain processing may stem from variable anatomy of SI in individual participants and, more importantly, the precise, localized somatotopic organization of SI (Bushnell et al., 1999). Our results cohere with this interpretation, as SI appeared only in analyses where there was less variability in the stimulation site (e.g., left-sided pain). Considering the highly somatotopic organization of SI, meta-analyses may need to focus on experiments of pain that target precisely the same physical locations of pain induction in order to reliably detect SI activation.

While there were subtle differences in which specific brain areas showed significant convergence across our main effect analyses, we found very few significant differences between experiments. The significant differences we did observe appeared to be driven by small differences in the extent of activation between experiments rather than qualitative differences in the underlying networks being recruited. Thus, it is possible that these effects reflect other sources of variability in the experimental paradigms, such as stimulation magnitude, stimulus duration, or variability in scanning protocol. It is important to note that in our analyses of differences in experimental paradigms (e.g., pain modality, pain location), we were limited to between-experiment contrasts, and meta-analyses of articles using within-subject contrasts would have a far greater ability to detect more nuanced differences between paradigms. Currently there were not enough experiments employing these designs to allow for such a meta-analysis. For this reason, future experiments using within-subject experimental designs are needed to further assess differences with greater sensitivity.

### 4.3. Limitations and future directions

Several limitations should be noted. As mentioned above, our findings delineate a convergent set of brain regions recruited across different experimental paradigms of pain, but an explicit meta-analysis of experiments using within-subject contrasts would better elucidate true differences in brain activation between paradigms. Given the limited number experiments that did explicitly test these differences within-subject, there is great potential for future experiments and meta-analyses to examine differences between experimental paradigms more conclusively. Similarly, because our meta-analysis relied on the published results of past articles, publication bias towards selectively publishing significant findings (and, consequently, difficulty in accounting for unpublished results) may have limited our findings (Müller et al., 2018). Additionally, it is possible that the wide range of image acquisition parameters in the experiments included may have limited the likelihood of observing some types of effects, particularly if detection sensitivity is realated to specialized acquisition parameters (e.g., brainstem, as mentioned in Sclocco et al., 2018). While we cannot exclude the possibility of other regions being more robustly involved in or associated with pain, open sharing of imaging data may allow for better detection of these regions using image-based meta-analytic methods (Salimi-Khorshidi et al., 2009) rather than solely using coordinate-based methods.

## 5. CONCLUSION

Convergent results demonstrate that SII, insula, ACC, and thalamus are consistently recruited by acute nociceptive stimuli in healthy subjects across many different experimental paradigms. In contrast, lateral PFC, M1, SMA, cerebellum, brainstem, SI, and the amygdala appear from the current meta-analysis to be more variably involved. Notably, we did not find strong evidence for preferential involvement of any of these brain areas in one specific experimental stimulus paradigm over another. Taken together, these findings suggest that acute pain induction in healthy volunteers consistently recruits a core brain network. If this network overlaps with that which is involved in clinical pain sensations, this “pain biomarker” may offer translational opportunities for fMRI in drug development by evaluation of analgesic efficacy and in clinical trials.

## ACKNOWLEDGEMENTS

Thank you to Zaixu Cui for his assistance in guidance of surface figure generation.

