## Supplementary Table 1 for "Convergent neural representations of acute nociceptive pain in healthy volunteers: A large-scale fMRI meta-analysis"

**Supplementary Table 1.** Experimental contributions to each significant cluster in main effect of all studies inducing pain (n = 200)

| Cluster | Experiments |
| --- | --- |
| Cluster 1: 14522 voxels (58, -24, 22) |  |
|  | Ando 2016 |
|  | Ariji 2018 |
|  | Asghar 2015 |
|  | Asghar 2016 |
|  | Atlas 2010 |
|  | Atlas 2014 |
|  | Baliki 2006 |
|  | Baliki 2009a |
|  | Baliki 2010 |
|  | Bar 2007 |
|  | Becker 2017 |
|  | Benson 2012 |
|  | Bingel 2007a |
|  | Bogdanov 2015 |
|  | Boland 2014 |
|  | Bouhassira 2013 |
|  | Boyle 2007 |
|  | Brinkmeyer 2010 |
|  | Brooks 2017 |
|  | Brugger 2011 |
|  | Brugger 2012 |
|  | Choi 2011 |
|  | Choi 2016 |
|  | Cleve 2017 |
|  | Coen 2008 |
|  | Coen 2009 |
|  | Coen 2011 |
|  | Cole 2006 |
|  | Cole 2010 |
|  | Corradi-Dell' Acqua 2011 |
|  | Davis 2016 |
|  | De la Fuente-Sandoval 2010 |
|  | De la Fuente-Sandoval 2012 |
|  | Dobek 2014 |
|  | Downar 2003 |
|  | Dube 2009 |
|  | Dunckley 2005a |
|  | Eisenblatter 2017 |
|  | Elsenbruch 2009 |
|  | Esser 2017 |
|  | Ettlin 2009 |
|  | Farmer 2013 |
|  | Farrell 2012 |
|  | Farrell 2014 |
|  | Fehse 2015 |
|  | Forkmann 2013 |
|  | Frankenstein 2001 |
|  | Freund 2007 |
|  | Freund 2009 |
|  | Gard 2012 |
|  | Geuze 2007 |
|  | Godinho 2012 |
|  | Gracely 2002 |
|  | Grant 2011 |
|  | Gu 2016 |
|  | Guleria 2017 |
|  | Gundel 2008 |
|  | Habig 2017 |
|  | Hahn 2013 |
|  | Hansen 2015 |
|  | Heckel_2011 |
|  | Hu 2015 |
|  | Iannilli 2008 |
|  | Ibinson 2013 |
|  | Jahn 2016 |
|  | Jensen 2015 |
|  | Kamping 2016 |
|  | Kattoor 2013 |
|  | Kim 2013b |
|  | Kobuch 2017 |
|  | Kobuch 2018 |
|  | Kong 2006 |
|  | Kong 2010 |
|  | Koyama 2005 |
|  | Kross 2011 |
|  | LaCesa 2014 |
|  | Ladabaum 2007 |
|  | Landgrebe 2008 |
|  | Lee 2008 |
|  | Lee 2015 |
|  | Leung 2016a |
|  | Lindstedt 2011 |
|  | Lloyd 2008 |
|  | Loggia 2012 |
|  | Loggia 2014 |
|  | Loken 2017 |
|  | Longo 2012 |
|  | Lopez-Sola 2010a |
|  | Lopez-Sola 2010b |
|  | Lu 2004 |
|  | Lui 2008 |
|  | Lutz 2013 |
|  | Lynn 2016 |
|  | Maeda 2011 |
|  | Maihofner 2004 |
|  | Maihofner 2005 |
|  | Maihofner 2006 |
|  | Maihofner 2011 |
|  | Mainero 2007 |
|  | Markl 2013 |
|  | Martin 2013 |
|  | Mayhew 2013 |
|  | Meier 2015 |
|  | Misra 2015 |
|  | Moana-Filho 2015 |
|  | Mobascher 2009a |
|  | Mobascher 2009b |
|  | Mobascher 2010a |
|  | Mobascher 2010b |
|  | Mochizuki 2007 |
|  | Mohr 2008 |
|  | Moisset 2010 |
|  | Morrison 2004 |
|  | Moulton 2011 |
|  | Moulton 2012 |
|  | Naglatzki 2012 |
|  | Nickel 2014 |
|  | Niddam 2002 |
|  | Obermann 2009 |
|  | Ochsner 2006 |
|  | Oertel 2008 |
|  | Oertel 2012 |
|  | Orenius 2017 |
|  | Oshiro 2007 |
|  | Oshiro 2009 |
|  | Pazmany 2017 |
|  | Peltz 2011 |
|  | Perini 2013 |
|  | Perlaki 2015 |
|  | Perrotta 2017 |
|  | Petschow 2016 |
|  | Piche 2010 |
|  | Pogatzki-Zahn 2010 |
|  | Pujol 2017 |
|  | Quiton 2014 |
|  | Roberts 2008 |
|  | Rodriguez-Raecke 2010 |
|  | Rosenberger 2009 |
|  | Rottmann 2010 |
|  | Roy 2009 |
|  | Rubio 2015 |
|  | Russo 2012 |
|  | Rutgen 2015 |
|  | Salomons 2015 |
|  | Scheef 2012 |
|  | Schenk 2017 |
|  | Schmahl 2006 |
|  | Schoell 2010 |
|  | Schulte 2016 |
|  | Schulz-Stubner 2004 |
|  | Seidel 2015 |
|  | Seifert 2007 |
|  | Seminowicz 2006 |
|  | Seminowicz 2007 |
|  | Sevel 2015 |
|  | Shelton 2012 |
|  | Shenoy 2011 |
|  | Shinozaki 2016 |
|  | Sinke 2016 |
|  | Sinke 2017 |
|  | Smith 2011 |
|  | Song 2006 |
|  | Sprenger 2015 |
|  | Sprenger 2018 |
|  | Stammler 2008 |
|  | Stankewitz 2010 |
|  | Starr 2009 |
|  | Stoeckel 2016 |
|  | Strigo 2013a |
|  | Strigo 2013b |
|  | Takahashi 2011 |
|  | Talmi 2009 |
|  | Tan 2015 |
|  | Tedeschi 2015 |
|  | Tessitore 2017 |
|  | Theysohn 2014 |
|  | Tseng 2010 |
|  | Tseng 2013 |
|  | Tseng 2015 |
|  | Tseng 2017 |
|  | Vachon-Presseau 2013 |
|  | van den Bosch 2013 |
|  | Vanhaudenhuyse 2009 |
|  | von Leupoldt 2008 |
|  | von Leupoldt 2009 |
|  | Wagner 2009 |
|  | Weiss 2008 |
|  | Wiech 2005 |
|  | Wiech 2006 |
|  | Wiech 2009 |
|  | Wiech 2010 |
|  | Winston 2014 |
|  | Woo 2015 |
|  | Yang 2012 |
|  | Yang 2018 |
|  | Yoshino 2010 |
|  | Youssef 2016 |
|  | Zeidan 2015 |
|  | Ziv 2010 |
| Cluster 2: 3181 voxels (6, 12, 38) |  |
|  | Ando 2016 |
|  | Asghar 2015 |
|  | Asghar 2016 |
|  | Atlas 2010 |
|  | Atlas 2014 |
|  | Baliki 2006 |
|  | Baliki 2009a |
|  | Baliki 2010 |
|  | Becker 2017 |
|  | Benson 2012 |
|  | Bogdanov 2015 |
|  | Boland 2014 |
|  | Brinkmeyer 2010 |
|  | Brooks 2017 |
|  | Brugger 2011 |
|  | Choi 2011 |
|  | Choi 2016 |
|  | Coen 2008 |
|  | Coen 2009 |
|  | Coen 2011 |
|  | Cole 2006 |
|  | Cole 2010 |
|  | Corradi-Dell' Acqua 2011 |
|  | Davis 2016 |
|  | De la Fuente-Sandoval 2010 |
|  | De la Fuente-Sandoval 2012 |
|  | Dobek 2014 |
|  | Downar 2003 |
|  | Dube 2009 |
|  | Dunckley 2005a |
|  | Eisenblatter 2017 |
|  | Esser 2017 |
|  | Ettlin 2009 |
|  | Farmer 2013 |
|  | Farrell 2012 |
|  | Farrell 2014 |
|  | Fehse 2015 |
|  | Ferris 2016 |
|  | Forkmann 2013 |
|  | Frankenstein 2001 |
|  | Freund 2009 |
|  | Gard 2012 |
|  | Geuze 2007 |
|  | Godinho 2012 |
|  | Gracely 2002 |
|  | Grant 2011 |
|  | Guleria 2017 |
|  | Gundel 2008 |
|  | Habig 2017 |
|  | Hansen 2015 |
|  | Hu 2015 |
|  | Iannilli 2008 |
|  | Ibinson 2013 |
|  | Jahn 2016 |
|  | Jensen 2015 |
|  | Kamping 2016 |
|  | Kobuch 2017 |
|  | Kobuch 2018 |
|  | Kong 2006 |
|  | Kong 2010 |
|  | Koyama 2005 |
|  | Kross 2011 |
|  | Ladabaum 2007 |
|  | Landgrebe 2008 |
|  | Lee 2008 |
|  | Loggia 2012 |
|  | Longo 2012 |
|  | Lopez-Sola 2010a |
|  | Lopez-Sola 2010b |
|  | Lu 2004 |
|  | Lui 2008 |
|  | Lutz 2013 |
|  | Lynn 2016 |
|  | Maeda 2011 |
|  | Maihofner 2006 |
|  | Maihofner 2011 |
|  | Mainero 2007 |
|  | Markl 2013 |
|  | Martin 2013 |
|  | Mayhew 2013 |
|  | Misra 2015 |
|  | Mobascher 2009a |
|  | Mobascher 2009b |
|  | Mobascher 2010a |
|  | Mobascher 2010b |
|  | Mochizuki 2007 |
|  | Mohr 2008 |
|  | Morrison 2004 |
|  | Moulton 2011 |
|  | Moulton 2012 |
|  | Naglatzki 2012 |
|  | Nickel 2014 |
|  | Niddam 2002 |
|  | Obermann 2009 |
|  | Ochsner 2006 |
|  | Oertel 2008 |
|  | Oertel 2012 |
|  | Oshiro 2007 |
|  | Oshiro 2009 |
|  | Peltz 2011 |
|  | Perlaki 2015 |
|  | Perrotta 2017 |
|  | Petschow 2016 |
|  | Piche 2010 |
|  | Pujol 2017 |
|  | Quiton 2014 |
|  | Roberts 2008 |
|  | Rodriguez-Raecke 2010 |
|  | Rottmann 2010 |
|  | Roy 2009 |
|  | Russo 2012 |
|  | Rutgen 2015 |
|  | Salomons 2015 |
|  | Scheef 2012 |
|  | Schenk 2017 |
|  | Schmahl 2006 |
|  | Schulz-Stubner 2004 |
|  | Seminowicz 2006 |
|  | Seminowicz 2007 |
|  | Sevel 2015 |
|  | Shelton 2012 |
|  | Shenoy 2011 |
|  | Shinozaki 2016 |
|  | Sinke 2016 |
|  | Sinke 2017 |
|  | Song 2006 |
|  | Sprenger 2015 |
|  | Sprenger 2018 |
|  | Stankewitz 2010 |
|  | Starr 2009 |
|  | Stoeckel 2016 |
|  | Strigo 2013b |
|  | Takahashi 2011 |
|  | Talmi 2009 |
|  | Tan 2015 |
|  | Tedeschi 2015 |
|  | Tessitore 2017 |
|  | Theysohn 2014 |
|  | Tseng 2013 |
|  | Tseng 2015 |
|  | Tseng 2017 |
|  | Vachon-Presseau 2013 |
|  | Vanhaudenhuyse 2009 |
|  | von Leupoldt 2008 |
|  | von Leupoldt 2009 |
|  | Wagner 2009 |
|  | Weiss 2008 |
|  | Wiech 2005 |
|  | Wiech 2010 |
|  | Winston 2014 |
|  | Woo 2015 |
|  | Yang 2012 |
|  | Yang 2018 |
|  | Yoshino 2010 |
|  | Youssef 2016 |
|  | Zeidan 2015 |
|  | Ziv 2010 |
| Cluster 3: 325 voxels (-32, -56, -34) |  |
|  | Ariji 2018 |
|  | Asghar 2015 |
|  | Bar 2007 |
|  | Benson 2012 |
|  | Bogdanov 2015 |
|  | Boland 2014 |
|  | Boyle 2007 |
|  | Brooks 2017 |
|  | Corradi-Dell' Acqua 2011 |
|  | Dobek 2014 |
|  | Dube 2009 |
|  | Farrell 2012 |
|  | Farrell 2014 |
|  | Gracely 2002 |
|  | Hahn 2013 |
|  | Hansen 2015 |
|  | Jahn 2016 |
|  | Kamping 2016 |
|  | Kim 2013b |
|  | Kong 2010 |
|  | Kross 2011 |
|  | Lee 2008 |
|  | Loggia 2014 |
|  | Lopez-Sola 2010a |
|  | Lu 2004 |
|  | Mobascher 2009a |
|  | Mobascher 2010a |
|  | Mobascher 2010b |
|  | Mohr 2008 |
|  | Nickel 2014 |
|  | Oshiro 2009 |
|  | Roy 2009 |
|  | Rutgen 2015 |
|  | Scheef 2012 |
|  | Shelton 2012 |
|  | Shenoy 2011 |
|  | Shinozaki 2016 |
|  | Song 2006 |
|  | Sprenger 2015 |
|  | Sprenger 2018 |
|  | Starr 2009 |
|  | Stoeckel 2016 |
|  | Strigo 2013b |
|  | Takahashi 2011 |
|  | Tedeschi 2015 |
|  | Weiss 2008 |
|  | Winston 2014 |
|  | Woo 2015 |
|  | Youssef 2016 |
| Cluster 4: 238 voxels (48, 4, 42) |  |
|  | Boyle 2007 |
|  | Brooks 2017 |
|  | Choi 2011 |
|  | Coen 2011 |
|  | Cole 2006 |
|  | Cole 2010 |
|  | Dube 2009 |
|  | Farmer 2013 |
|  | Farrell 2012 |
|  | Farrell 2014 |
|  | Freund 2007 |
|  | Gard 2012 |
|  | Geuze 2007 |
|  | Iannilli 2008 |
|  | Kong 2010 |
|  | Kross 2011 |
|  | Ladabaum 2007 |
|  | Landgrebe 2008 |
|  | Loggia 2012 |
|  | Longo 2012 |
|  | Lopez-Sola 2010a |
|  | Maihofner 2005 |
|  | Maihofner 2006 |
|  | Maihofner 2011 |
|  | Mayhew 2013 |
|  | Misra 2015 |
|  | Moisset 2010 |
|  | Moulton 2011 |
|  | Ochsner 2006 |
|  | Oshiro 2009 |
|  | Peltz 2011 |
|  | Piche 2010 |
|  | Quiton 2014 |
|  | Russo 2012 |
|  | Salomons 2015 |
|  | Scheef 2012 |
|  | Schmahl 2006 |
|  | Shinozaki 2016 |
|  | Stoeckel 2016 |
|  | Takahashi 2011 |
|  | Tseng 2010 |
|  | Tseng 2015 |
|  | von Leupoldt 2009 |
|  | Wagner 2009 |
|  | Winston 2014 |
|  | Ziv 2010 |

*Note*: Cluster identification on the first column indicates cluster number, number of voxels in cluster, and peak coordinate of cluster.
